# Secondary growth in the tomato stem is suppressed by the flowering regulators FUL2 and STM3

**DOI:** 10.64898/2026.11.04.763880

**Authors:** Kailash Pandey, Amponsah Preko Appiah, Robin van Velzen, Iris E. Zahn, Ellora Basu, Gerco C. Angenent, Peter J. Deuss, Marian Bemer

**Affiliations:** Laboratory of Molecular Biology, Wageningen University & Research, Droevendaalsesteeg 1, 6708 PB Wageningen, the Netherlands; Biosystematics group, Wageningen University & Research, Droevendaalsesteeg 1, 6708 PB Wageningen, the Netherlands; Business Unit Bioscience, Wageningen University & Research, Droevendaalsesteeg 1, 6708 PB Wageningen, The Netherlands; Department of Chemical Engineering, Engineering and Technology Institute Groningen (ENTEG), University of Groningen, Nijenborgh 3, Groningen 9747 AG, the Netherlands; Leiden University, Institute of Biology Leiden (IBL), Plant Sciences, Sylviusweg 72, 2333 BE Leiden, The Netherlands

## Abstract

Herbaceous eudicot crops such as tomato accumulate substantial amounts of secondary xylem in their stems, particularly in the basal part. As plants progress from the vegetative phase to flowering and fruit development, secondary growth continues to provide mechanical support and facilitate transport. To uncover how secondary growth is regulated in the tomato stem, we used laser capture microdissection RNA-seq to establish tissue-specific transcriptomic profiles. This revealed that key regulators associated with wood formation in poplar are also acting in tomato. Furthermore, we found that the flowering regulators *FUL2* and *STM3* are substantially expressed in xylem tissue. Subsequent functional analysis identified FUL2 and STM3 as key repressors of secondary growth, with *ful2* or *stm3* knock-out mutants exhibiting strongly increased secondary growth in older tomato stems. Notably, a higher order mutant in which both *ful2* and *stm3* were knocked out displays a similar non-additive phenotype, indicating, together with yeast two-hybrid data showing physical interaction, that FUL2 and STM3 act together in a complex. The expression levels of both genes remain low until after the floral transition but strongly rise during subsequent stem development. Transcriptomic analyses in *ful2*, *stm3,* and wild-type stems at stages before and after detectable anatomical differences, showed that the FUL2-STM3 complex represses distinct sets of *NAC* and *MYB* candidate regulators associated with xylem development. Interestingly, FUL2 and STM3 control these regulators in a stage- and position-dependent way, with candidates showing upregulation only at later stages in mid-height stems being already upregulated at earlier stages in basal stem regions. Our findings provide a tissue-resolved transcriptomic resource for tomato stems and reveal a role for FUL2 and STM3 in limiting secondary xylem accumulation as stem development progresses.

## INTRODUCTION

Plants extend their longitudinal axes through primary growth and increase in width during secondary growth. Primary growth is driven by apical meristems, whereas secondary growth is sustained by the vascular cambium, a lateral meristem that produces secondary xylem and secondary phloem (De Rybel et al., 2016; Shi et al., 2019; Wybouw et al., 2024). These vascular tissues enable long-distance transport of water, nutrients and photosynthates, while secondary xylem also provides mechanical support required throughout the plant lifespan. Secondary xylem, or wood, is a major component of terrestrial biomass and an important renewable biological resource (Bar-On et al., 2018; Wang et al., 2021). Moreover, xylem hydraulic function appears to be directly related to stress resilience and recovery (Brodribb et al., 2010; Hendriks et al., 2026; Lens et al., 2022; Rowland et al., 2015). Secondary growth is therefore central to plant development and structural resilience and contributes to the lignocellulosic biomass that accumulates in stems.

Secondary growth is not limited to trees. Herbaceous dicots, including the model species Arabidopsis and tomato, can establish a vascular cambium and produce substantial secondary xylem (Chaffey et al., 2002; Coaker et al., 2002; Lens et al., 2012; Nieminen et al., 2004). Its extent nevertheless varies widely among species, between organs, and across developmental stages, indicating that radial growth is a strongly regulated developmental process rather than simply a consequence of prolonged growth (Spicer & Groover, 2010; Tomescu & Groover, 2019).

With the progression of secondary growth, the vascular cambium produces more secondary xylem than secondary phloem, leading to an increasing proportion of xylem within the vascular cylinder (Carlsbecker & Augstein, 2021; Tomescu & Groover, 2019). Secondary xylem formation then proceeds through a coordinated developmental sequence. The xylem side derivatives of the cambium undergo cell expansion, secondary cell wall deposition and lignification as they differentiate into specialized xylem cell types (Nieminen et al., 2004; Ye & Zhong, 2015). Because these steps are sequential and partly separately controlled, the amount of xylem formed reflects the combined output of cambial proliferation, xylem differentiation and secondary cell wall formation, and can be altered at any one of them. The cambial region itself is spatially organized into proximal, central and distal domains, corresponding broadly to the xylem facing side, bifacial stem cell region and phloem facing side of the cambium, respectively. In Arabidopsis, overlapping expression of *PHLOEM INTERCALATED WITH XYLEM (PXY)* and *WUSCHEL-RELATED HOMEOBOX4 (WOX4)* marks the central and partially proximal domains, whereas *SUPPRESSOR OF MAX2 1-LIKE5 (SMXL5)* marks the central and partially distal domains (Shi et al., 2019). Cambial proliferation is regulated mainly by the PXY-WOX4 regulatory module (Etchells & Turner, 2010; Hirakawa et al., 2010; Wang et al., 2019), while HD-ZIP III transcription factors (TFs) control vascular patterning and xylem identity determination (Bishopp et al., 2011; Carlsbecker et al., 2010; Kondo et al., 2014; Ramachandran et al., 2017). Following xylem specification, NAC- and MYB-domain TFs coordinate xylem differentiation and secondary cell wall formation. VASCULAR-RELATED NAC-DOMAIN6 (VND6) and VND7 promote vessel differentiation (Lehmann & Schneider, 2025; Yamaguchi et al., 2010, 2011), and NAC SECONDARY WALL THICKENING PROMOTING FACTOR 1 (NST1) and NST3 regulate fiber differentiation and secondary cell wall formation (Mitsuda et al., 2007; Yamaguchi et al., 2010; Zhong et al., 2007). Downstream of these NAC regulators, MYB46 and MYB83 regulate secondary cell wall related program, while MYB58 and MYB63 activate lignin biosynthesis (Ko et al., 2012; McCarthy et al., 2009; Zhou et al., 2009). The regulation of secondary xylem formation is therefore relatively well described, but the developmental cues that determine when substantial secondary xylem production is initiated remain much less understood.

Secondary growth is itself a developmental feature and is regulated in coordination with other developmental processes. In the Arabidopsis hypocotyl, floral induction leads to a modification of secondary growth regulation. Flowering promotes the transition from a phase of proportional xylem and phloem production to the xylem expansion phase through a shoot-derived mobile gibberellin signal (Ragni et al., 2011; Sibout et al., 2008). Soon thereafter, cambial activity decreases and the cambium undergoes senescence, resulting in a woody hypocotyl that supports the inflorescence stem that starts to elongate (Ben-Targem et al., 2021). How this coupling between flowering and secondary growth regulation functions in the inflorescence stem, which is formed after the floral transition and whose cambium does not undergo senescence upon flowering, is not understood.

A direct genetic link between flowering time regulators and secondary growth has been established in the Arabidopsis inflorescence stem. The flowering time genes *FRUITFULL* (*FUL*) and *SUPPRESSOR OF OVEREXPRESSION OF CONSTANS 1* (*SOC1*) suppress secondary growth, and the *ful soc1* double mutant shows extensive secondary xylem formation and a perennial growth habit (Melzer et al., 2008). The MADS-domain transcription factors FUL and SOC1, which can form heterodimers and heterotetramers (de Folter et al., 2005; Thoris et al., 2024), repress *AT-HOOK MOTIF CONTAINING NUCLEAR LOCALIZED 15* (*AHL15*), which encodes a transcription factor that positively regulates cambial activity and secondary growth (Karami et al., 2020; Rahimi et al., 2022).Thus, interestingly, the induction of flowering is in Arabidopsis coupled to the repression of inflorescence stem secondary growth via the FUL-SOC1-AHL15 module.

So far, the genetic regulation of stem secondary growth has been primarily investigated in the tree species poplar and the short-living herbaceous species Arabidopsis. Other herbaceous species, such as the important crop tomato, produce substantially more wood than Arabidopsis, but it is currently unknown how secondary growth is regulated in tomato. Tomato is an extensively studied model species for understanding flowering time regulation (Jiang et al., 2025; Lifschitz et al., 2006; MacAlister et al., 2012), inflorescence architecture (Lippman et al., 2008; Park et al., 2012; Soyk et al., 2017), and fruit development (Li et al., 2026; The Tomato Genome Consortium, 2012; Wang & de Maagd, 2025; Zhang et al., 2024). In contrast, the molecular regulation of tomato stem development, particularly secondary growth, has hardly been studied.

Interestingly, a regulatory link between flowering and secondary growth has also been described in tomato, albeit different from Arabidopsis. In tomato, elevated activity of the florigen SINGLE FLOWER TRUSS (SFT), a key activator of the reproductive transition, accelerates secondary cell wall biogenesis, while *35S:FUL2* lines exhibit slender stems with premature lignification (Shalit-Kaneh et al., 2019). These findings show that increased activity of flowering-associated regulators can modify tomato vascular development. Similar to Arabidopsis, the tomato FUL-like proteins FUL2 and MBP20, and SOC1-like proteins TOMATO MADS 3 (TM3) and SISTER OF TM3 (STM3) interact with each other in MADS-box complexes to induce the transition from vegetative meristem to reproductive meristem by binding to CArG-boxes of target genes such as *APETALA 2* (*AP2*) and cytokinin oxidase genes (Jiang et al., 2022, 2025; Zahn et al., 2023). However, how secondary growth in tomato is regulated and what the endogenous contributions of flowering genes such as *SFT, FUL2, and TM3/STM3* to tomato stem secondary growth are, have remained unclear.

Here, we investigated secondary growth in the tomato stem and performed spatial transcriptomics to gain insight into the genes that are predominantly expressed in the cambium and xylem. This analysis revealed that the key genes that regulate secondary growth in the poplar tree are also acting in the tomato stem, indicating that the core regulatory framework for secondary xylem formation is shared between herbaceous and woody species. Moreover, the spatial transcriptomics data also suggested functions for co-orthologs of the Arabidopsis genes *FUL* and *SOC1*. We further studied CRISPR/Cas9 single and higher order mutants of tomato *FUL*- and *SOC1*-like genes across successive stem development stages and identified FUL2 and STM3 as key repressors of secondary growth. Their expression in the stem rises after the floral transition and impacts secondary growth when the stem reaches a more advanced developmental state. Transcriptome profiling at two stages after transition to flowering, before and after the mutant phenotype becomes apparent, revealed that the regulatory output of FUL2 and STM3 changes with stem development. Together, FUL2 and STM3 repress genes encoding NAC- and MYB-domain TFs associated with xylem differentiation and secondary cell wall formation. The resulting changes in secondary growth also alter lignin content in older stems. Our findings reveal a spatially and temporally structured function of flowering related MADS-domain transcription factors in repressing secondary growth in the tomato stem.

## RESULTS

### Tomato stem secondary growth varies across temporal and spatial developmental axes

In many dicot stems, secondary growth is driven by a ring of vascular cambium formed through the connection of intrafascicular cambium within the primary vascular bundles and interfascicular cambium formed between the bundles (Altamura et al., 2001; Nieminen et al., 2015). As secondary growth progresses, the vascular cambium produces secondary xylem towards the inside and secondary phloem towards the outside (Figure 1A). The extent of secondary xylem formation varies with both plant age and position along the stem (Baucher et al., 2007; Chao et al., 2019; Teichmann et al., 2008). Consistent with this developmental progression, younger tomato plants show relatively limited secondary xylem accumulation, whereas substantially more secondary xylem is present at later stages (Figure 1B). Secondary growth also differs between positions within the same plant. At 120 days after germination (DAG), the basal stem shows substantially greater secondary xylem accumulation than the mid-height region (Figure 1B). Thus, chronological age alone does not describe the secondary growth status of a particular stem region. Here, we use the term local secondary growth state to describe the relative progression of secondary growth at a given stem position, as reflected by secondary xylem accumulation. The term describes relative progression rather than a terminal state, as stems of indeterminate tomato plants continue to undergo secondary growth throughout development.

**Figure 1.**
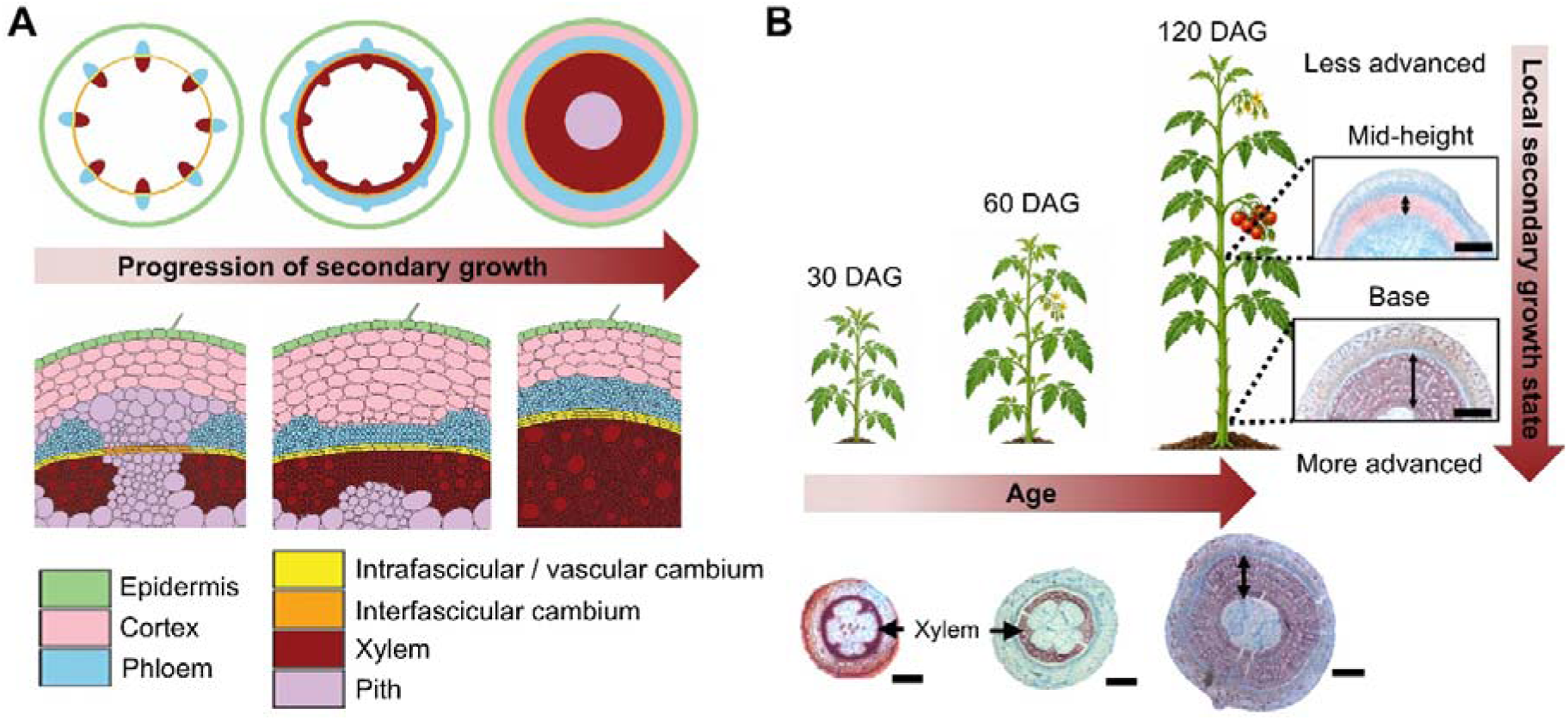
Stem secondary growth varies with plant age and position along the stem. **(A)** Schematic representation of progression of secondary growth in a dicot stem, showing the formation of continuous vascular cambium and the production of secondary xylem and secondary phloem. **(B)** Schematics of tomato plants with representative cross-sections illustrating secondary xylem accumulation across plant ages and at different stem positions. Double headed arrows indicate the radial extent of secondary xylem. DAG, days after germination. Scale bars = 2 mm.

### LCM RNA-seq defines tissue-associated transcriptional profiles during tomato secondary growth

To establish a spatial transcriptional framework for the tomato stem, we performed laser capture microdissection followed by RNA sequencing (LCM RNA-seq). We isolated four anatomically defined tissue fractions: cambium, xylem, pooled phloem+cortex, and pooled cortex+epidermis (Figure 2A). Samples were collected from the mid-height region of 60-day old plants, where an active cambium and developing xylem and phloem are evident (Figure 2A). To evaluate transcriptome coverage and reproducibility across the LCM samples, we performed principal component analysis (PCA). Biological replicates showed consistent transcript abundances and separated according to tissue fraction, while commonly used tomato reference genes showed stable expression across samples (Bokhale et al., 2020; González-Aguilera et al., 2016) (Supplemental Figures 1A-D; Supplemental Data Set 1). We next examined the tissue-associated transcriptional signatures using differential expression analysis followed by MapMan based gene set enrichment analysis (GSEA) (BH adjusted *p*<0.1) (Subramanian et al., 2005; Thimm et al., 2004; Usadel et al., 2005). The cambial sample was enriched for categories associated with chromatin organization, microtubule network, leucine rich repeat receptor-like kinase (LRR-RLK) signaling, and HD-ZIP III associated transcriptional regulation, whereas categories related to hormone repression, stress, transport, and metabolism were depleted (Supplemental Figure 1E). In xylem, categories associated with phenylalanine-based metabolism, lignin polymerization and cellulose synthesis were enriched, whereas photosynthesis, redox related processes, chromatin organization, and LRR-RLK signaling were negatively enriched (Supplemental Figure 1F).

**Figure 2.**
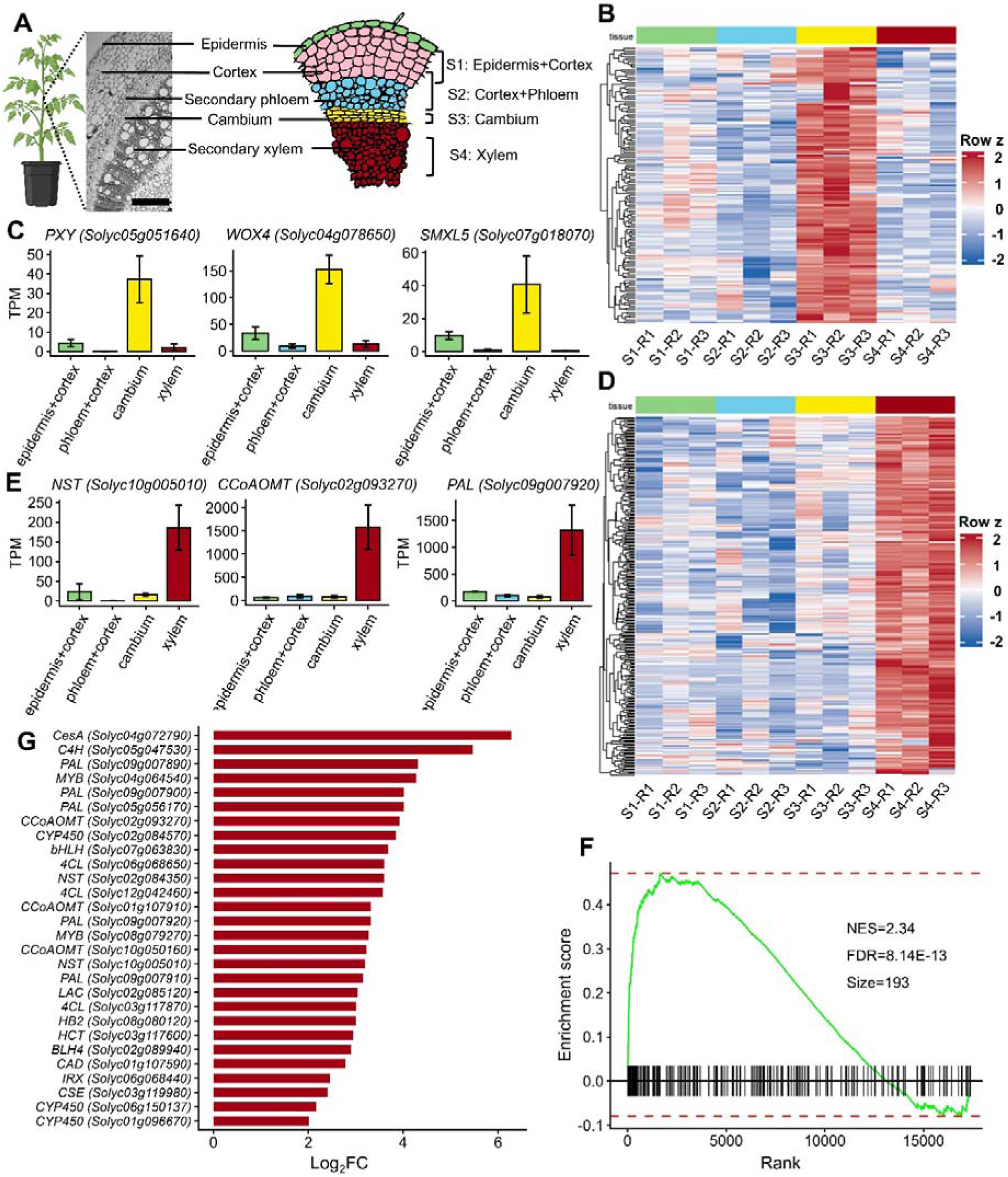
Tissue resolved transcriptomic profiling of tomato stem using LCM RNA-seq. **(A)** Graphical representation of the mid-height stem region of a 60-day old tomato plant and the tissues collected for LCM RNA-seq. A Steedman’s wax embedded stem sample was sectioned into 12 µm transverse sections, with the secondary xylem, cambium, phloem, cortex and epidermis labelled. Scale bars, 400 µm. The schematic on the right indicates the four dissected tissue fractions: S1, epidermis and cortex (green and pink); S2, cortex and phloem (blue); S3, cambium (yellow); and S4, secondary xylem (red). **(B)** Heatmap of VST normalized expression levels of cambium-specific genes across tissue types. A total of 147 cambium-specific genes were identified with enrichment ratio (ER) > 3. **(C)** TPM normalized expression of cambium-specific genes that are orthologs of marker genes from Arabidopsis: *PXY* and *WOX4* (central and proximal domains of the cambium) and *SMXL5* (distal domain of the cambium). Bars show mean TPM values ± SEM. **(D)** Heatmap of VST normalized expression levels of xylem-specific genes across tissue types. A total of 322 xylem-specific genes were identified with enrichment ratio (ER) > 3. **(E)** TPM normalized expression of xylem-specific genes that are orthologs of key xylem regulatory genes from Arabidopsis. Bars show mean TPM values ± SEM. **(F)** Gene set enrichment analysis (GSEA) plot of tomato orthologs of xylem-specific genes and key genes regulating secondary cell wall biogenesis and lignin biosynthesis in poplar, in the xylem vs rest LCM differential expression analysis. The curve shows the running enrichment score; black lines indicate the positions of gene-set members in the ranked list, and dotted lines mark the maximum and minimum enrichment scores. **(G)** Representative tomato orthologs among the leading-edge genes in (F), showing their shrunken log_2_FC values from the xylem vs rest LCM differential expression analysis.

To further resolve the tissue-specific expression, we calculated an enrichment ratio (ER) by comparing transcript abundance in each target tissue with the highest abundance observed among the remaining fractions. We defined genes as tissue specific if they met all of the following criteria: FDR < 0.05, ER > 3, shrunken log2 fold change > 1, mean TPM > 2, and raw counts ≥10 in at least two biological replicates. This identified 147 cambium-specific genes (Figure 2B; Supplemental Data Set 1). Among these are the tomato orthologs of the established Arabidopsis cambium regulators *PXY* (*Solyc05g051640*), *WOX4* (*Solyc04g078650*) and *SMXL5* (*Solyc07g018070*) (Figure 2C).

Similarly, we identified 322 xylem-specific genes (Figure 2D; Supplemental Data Set 1). These included tomato orthologs of established regulators of xylem differentiation and secondary wall formation, including *NAC SECONDARY WALL THICKENING PROMOTING FACTOR* (*NST*, *Solyc10g005010*), and genes encoding enzymes involved in lignin biosynthesis, such as caffeoyl-CoA 3-O-methyltransferase (*CCoAOMT, Solyc02g093270*), and phenylalanine ammonia lyase *(PAL, Solyc09g007920)* (Figure 2E). The recovery of these established cambial and xylem regulators independently supports the vascular domains resolved by the LCM. Furthermore, it indicates conservation of core cambial and xylem regulatory features between Arabidopsis and tomato.

Because tomato is an herbaceous eudicot that undergoes substantial secondary growth, we next asked whether broader transcriptional signatures associated with secondary xylem and wood formation are conserved between poplar and tomato. Using tomato orthologues of poplar xylem specific transcription factors and key genes involved in secondary cell wall and lignin biosynthesis (Supplemental Data Set 1), we observed strong enrichment of the poplar xylem signature within the tomato LCM sample (Figure 2F). The enriched genes encode amongst others MYB, NAC and bHLH transcription factors associated with secondary cell wall formation, different IRREGULAR XYLEM (IRX) factors, and core components of lignin and cellulose biosynthesis, including PAL, CCoAOMT, cinnamyl alcohol dehydrogenase (CAD), cinnamate 4-hydroxylase (C4H), 4-Coumarate: CoA ligase (4CL), cellulose synthase (CesA), caffeoyl shikimate esterase (CSE), and hydroxycinnamoyl transferase (HCT) (Figure 2G).

Altogether, these analyses show conservation of major cambium and xylem-related expression signatures across herbaceous and woody eudicots. This tissue-resolved dataset establishes a useful resource for studying stem development and identifying regulators associated with secondary growth in tomato.

### *FUL2* and *STM3* show tissue-associated expression in the LCM dataset

Given that the flowering time genes *FUL* and *SOC1* are important repressors of secondary growth in Arabidopsis, we subsequently focused on known tomato flowering genes to determine whether there is a link between the flowering time and secondary growth pathways in tomato. To this end, we composed a set of floral induction genes based on literature (Supplemental Table 1) and searched our LCM data for genes that are specifically expressed in cambium and/or xylem. We determined the tissue-associated expression scores using enrichment ratio (ER > 3), and depletion ratio (DR > 3) to measure the depletion from the epidermis+cortex sample. Interestingly, none of the flowering genes showed preferential association with the cambium, and only *FUL2* showed xylem-associated expression (Supplemental Figure 2). Of the other tomato flowering genes, expression was either not detected, very weak (TPM < 1 in all dissected tissue domains), or non-specific (Supplemental Figure 2). It is worth noting that *SINGLE FLOWER TRUSS (SFT)* transcripts were not detected in our LCM dataset, whereas *SFT* stem expression was reported by Shalit-Kaneh et al. (2019). Interestingly, although *STM3*, a tomato *SOC1* co-ortholog, was not specifically expressed in cambium or xylem, it was highly expressed in all inner vascular tissues (cambium, xylem, phloem/cortex). This indicates that the FUL-SOC1 module that exists in Arabidopsis may also act in tomato. Therefore, we focused on the *FUL*-like and *SOC1*-like genes in the LCM data. The tomato genome contains four *FUL-*like genes, *FUL1, FUL2, MADS-BOX PROTEIN 10* (*MBP10*) and *MBP20* (Maheepala et al., 2019), and three *SOC1*-like genes, *TOMATO MADS 3* (*TM3*), *SISTER OF TM3* (*STM3*) and *MBP23* (Zahn et al., 2023).

Among the *FUL*-like genes, *MBP20* was not detected, while *MBP10* was expressed at low levels in cambium and xylem. *FUL1* showed higher expression in the epidermis+cortex samples, whereas *FUL2* showed its highest expression in xylem (Figures 3A and 3B). Similarly, among the *SOC1*-like genes, *MBP23* was not detected, *TM3* was preferentially associated with the phloem+cortex, and *STM3* was highly expressed across the inner vascular tissues (Figures 3A and 3B). Thus, while our LCM transcriptome analysis does not indicate a general role for flowering genes in secondary growth, it suggests a conserved role for *FUL* and *SOC1* orthologs as regulators of both flowering time and secondary growth in tomato. Although we did not identify transcripts of the key flowering regulator *SFT*, the fact that *SFT* encodes the “florigen” protein that can travel through the phloem (Corbesier et al., 2007; Lifschitz et al., 2006) leaves open the possibility that it is an enhancer of secondary cell wall biogenesis in the stem, as suggested by the work of Shalit-Kaneh et al. (2019).

**Figure 3.**
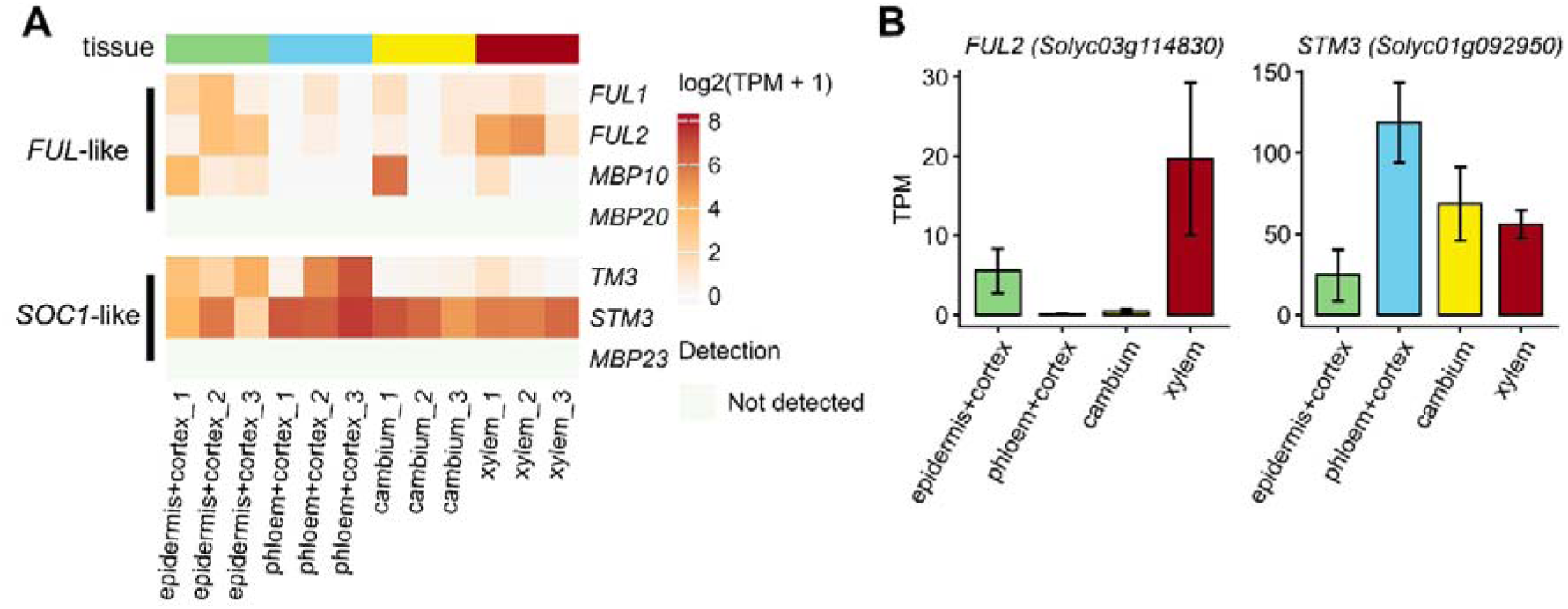
*FUL*- and *SOC1*-like genes show distinct expression patterns across stem tissues. **(A)** Expression profiles of *FUL*- and *SOC1*-like genes across four LCM samples. Expression is shown as log_2_(TPM+1). **(B)** Transcript abundance of *FUL2* and *STM3* across the four dissected samples. Values are shown as TPM; bars represent mean ± SEM.

### FUL2 and STM3 suppress secondary growth during stem development

To determine whether *FUL*- and *SOC1*-like genes are key regulators of stem secondary growth in tomato, we used CRISPR/Cas9 mutant lines previously characterized for their flowering time phenotypes (Jiang et al., 2022; Zahn et al., 2023). We first examined secondary growth at the basal internode of 120 DAG *ful2 mbp20 tm3 stm3* quadruple mutants (hereafter, *quad ful soc1*). At 120 DAG, plants of all genotypes were flowering. The quadruple mutant showed a marked increase in secondary xylem formation compared to the WT (Figure 4A), with the xylem area being doubled compared to WT. This reveals that *FUL*- and *SOC1*-like genes indeed contribute to the regulation of tomato secondary growth.

**Figure 4.**
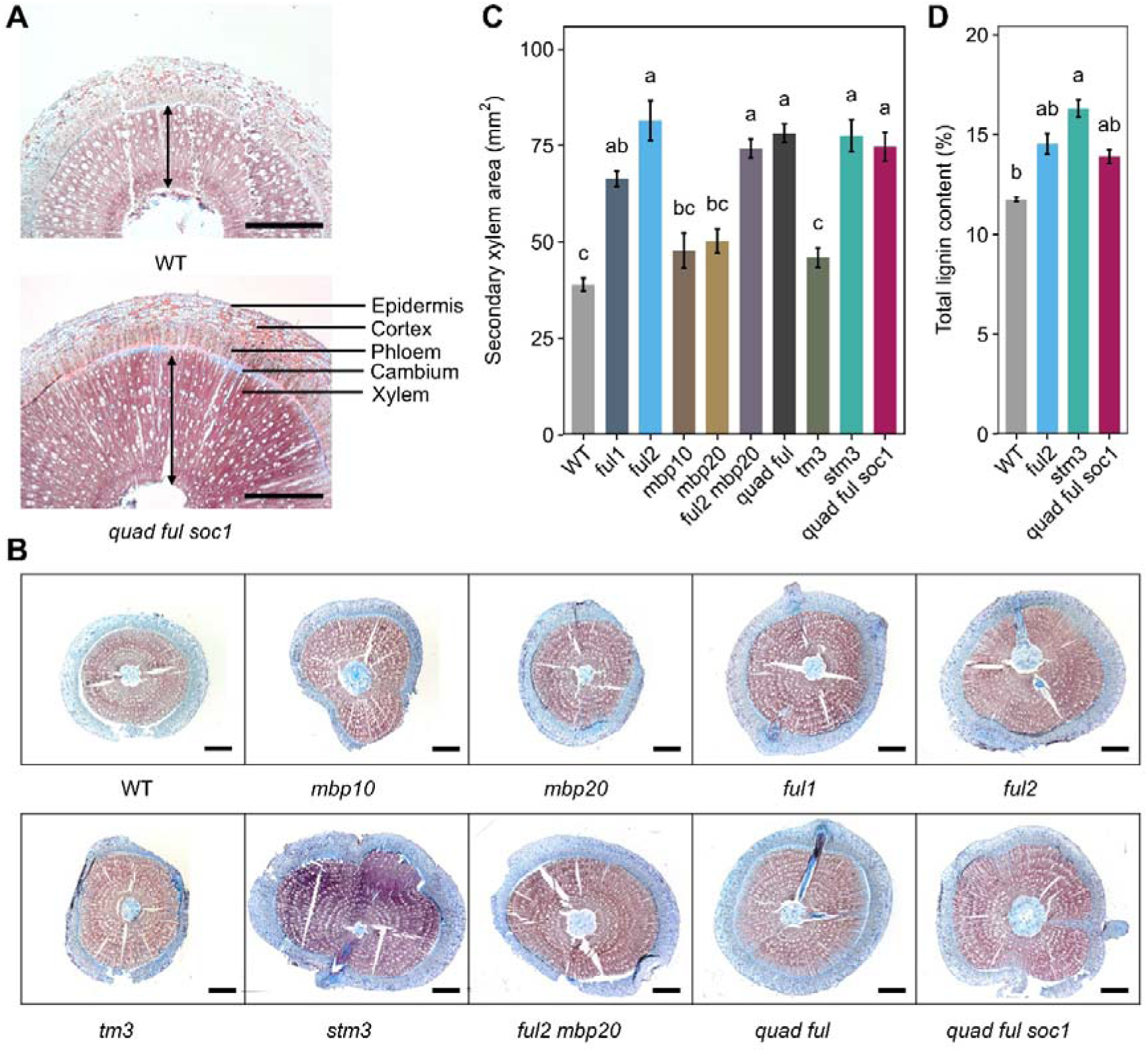
*FUL*- and *SOC1*-like genes suppress secondary growth in tomato stem. **(A)** Representative stem cross-sections of 120 DAG WT and *quad ful soc1* mutant plants. The quadruple mutant showed increased secondary xylem formation compared with WT. Black arrows show the width of secondary xylem. Scale bars, 2 mm. **(B)** Representative stem cross-sections of *FUL*- and *SOC1*-like single and higher order mutants at 120 DAG. **(C)** Quantification of secondary xylem area in 120 DAG plants. *ful2* and *stm3* showed the largest increases in stem area and secondary xylem area, which were not further increased in the higher order mutants. **(D)** Total lignin content in the bottom half of stem of the 120 DAG WT and mutant plants. Lignin content is expressed as a percentage on an extractive-free basis. *stm3* showed the largest increase in lignin content, whereas *ful2* and the *quad ful soc1* mutants showed modest increases. Statistical differences were determined by one-way ANOVA followed by Tukey’s HSD test (*p* < 0.01). Values are shown as mean ± SE (n = 4). Different letters indicate statistically significant differences.

We next phenotyped the basal internode of 120 DAG different single and higher order mutants to determine the contribution of individual genes. Among the single mutants, *ful2* and *stm3* showed the strongest increase in secondary xylem area compared to WT, and the phenotype was not further enhanced in higher order mutants in which *FUL2* and/or *STM3* were mutated (Figures 4B and 4C). Stem cross-sectional area showed a similar pattern, while xylem:phloem ratios were slightly more increased in *quad ful soc1* mutants than in both single mutants (Supplemental Figures 3A and 3B). The near absence of an additive phenotype in *quad ful soc1* mutants suggests that FUL2 and STM3 act in the same pathway and that the role of MBP20 and TM3 in the regulation of secondary xylem formation is very limited or absent. Given that the FUL2 and STM3 proteins can physically interact (Jiang et al., 2022; Zahn et al., 2023), this occurs most likely as partners in the same protein complex. Altogether, our results indicate that *FUL2* and *STM3* are the major *FUL*- and *SOC1*-like genes that are each required to suppress secondary growth in the tomato stem.

To check if the increase in secondary xylem formation was accompanied by changes in the cell-wall composition, we analyzed the biochemical composition of the basal half of the stem (Supplementary Figures 3C-G). *stm3* showed the largest increase in total lignin content per gram dry weight, from 11.8% in WT to 16.3%, a 38.6% relative increase in *stm3* stems. *ful2* and *quad ful soc1* showed intermediate increases in lignin levels, corresponding to numerical increases of 23.6% and 18.2% relative increase, respectively (Figure 4D). However, total carbohydrate content remained comparable to WT (Supplementary Figure 3E).

To understand whether the enhanced secondary growth phenotype is due to early acceleration of secondary growth or develops later during stem maturation, we checked the mutants at earlier developmental stages. At 30 DAG, secondary xylem area was comparable between the mutants and WT, although the *quad ful soc1* showed a slight increase (Supplemental Figures 4A-C). At 60 DAG, secondary xylem area remained similar to WT (Supplemental Figures 4D-F). We next checked the mid-height region of the 120 DAG plants, which is undergoing active secondary growth, but much less pronounced than at the base (Figure 1). At this position, *ful2*, *stm3* and higher order mutants already showed an increased secondary xylem-to-phloem ratio (Supplemental Figure 5). Therefore, although the enhanced xylem formation is only evident in the basal part of 120 DAG stems, altered allocation toward xylem is already apparent in the upper developing stem regions. These results suggest that the phenotype progressively develops during stem development, further supporting the major roles of *FUL2* and *STM3* in regulating the balance of secondary growth in the tomato stem.

### *FUL2* and *STM3* show overlapping expression patterns in the stem

To investigate the temporal and spatial gene activity of *FUL2* and *STM3* during plant development, we performed qRT-PCR experiments at different stages and positions. At 15 and 30 DAG, the transcript levels of both genes were rather low, while they were much higher at 60 and 120 DAG (Figures 5A and 5B). Under our growth conditions, the floral transition, defined as doming of the SAM, occurs at approximately 21 DAG. Thus, 15 DAG precedes and 30 DAG follows this transition. The weak expression at both stages suggests that the later increase in *FUL2* and *STM3* expression is not an immediate consequence of floral transition. Instead, their expression increases during subsequent stem development and remains broadly maintained at later stages. At 60 and 120 DAG, the expression also showed little variation along the height of the stem (Figures 5A and 5B). *TM3* has a broadly similar developmental expression pattern, albeit at lower levels, whereas *MBP20* was expressed at very low levels across the examined samples (Supplemental Figure 6). Thus, it is not surprising that the phenotype of the *quad ful soc1* mutant is similar to that of the *ful2* and *stm3* single mutants.

**Figure 5.**
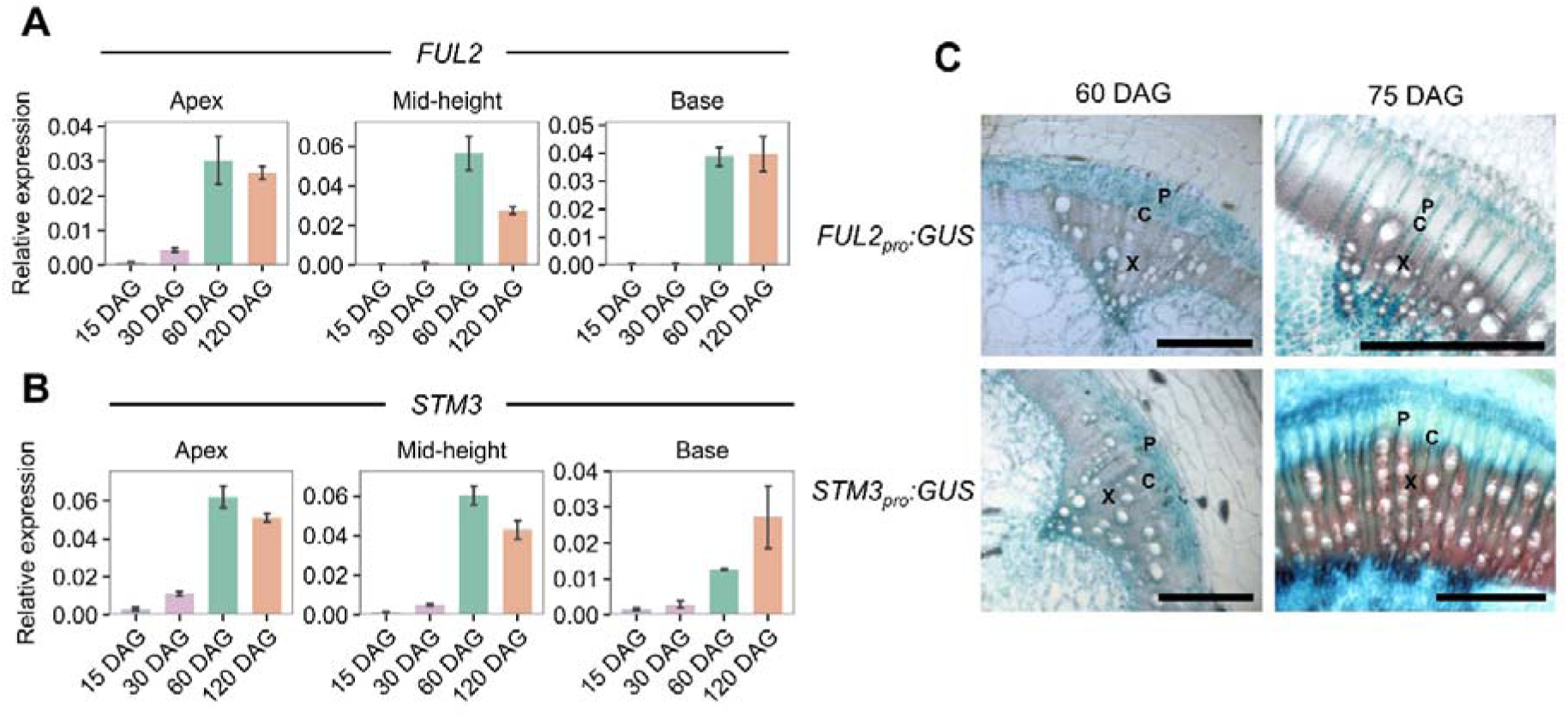
Spatial expression patterns of *FUL2* and *STM3* at different stages of stem maturation. **(A, B)** Expression of *FUL2* (A) and *STM3* (B) at 15, 30, 60 and 120 DAG across different positions along the stem height. Expression was relatively low at 15 and 30 DAG and increased at 60 DAG and 120 DAG with limited variation at later developmental stages. **(C)** Expression pattern of *FUL2_pro_:GUS* and *STM3_pro_:GUS* at 60 DAG and 75 DAG. The reporter activity was detected across the vascular tissues including secondary xylem, cambial zone and phloem. At 75 DAG, expression was prominent in ray cells across the secondary vascular tissues. P, phloem; C, Cambial zone; X, xylem. Scale bars, 500 µm.

We examined the spatial expression pattern of *FUL2* and *STM3* in more detail using promoter-reporter lines and found largely overlapping expression patterns (Figure 5C). At 60 DAG, GUS reporter activity was detected across the vascular tissues, including secondary xylem, cambial zone and phloem for both the *FUL2_pro_:GUS* and *STM3_pro_:GUS* lines. While this was for *STM3* completely in line with the LCM data, *FUL2_pro_:GUS* showed a broader pattern than expected, with also GUS signal in the cambium. This may be explained by the fact that the LCM data was derived from mid-height sections or by a discrepancy between the activity of the used promoter fragment and the actual transcript abundance. Interestingly, at 75 DAG, the expression remained associated with the vascular tissue but became prominent in the ray cells extending across the secondary vascular tissues, indicating a change in spatial promoter activity. It appears that in particular *FUL2* gets restricted to the xylem and ray cells at this stage. Ray cells form radial connections across the secondary vascular tissues and are involved in transport between the xylem and phloem, placing them in a suitable position to participate in radial signaling during stem development (Hunziker & Greb, 2024; Pfautsch, Hölttä, et al., 2015; Spicer, 2014). These experiments revealed an additional layer of complexity, with the spatial activity of *FUL2* and *STM3* also shifting in location based on developmental stage.

### FUL2 and STM3 regulate a distinct set of genes at 60 DAG

We performed RNA-seq transcriptome analysis with stem samples of *ful2, stm3* and WT at 60 and 120 DAG, before and after the occurrence of phenotypic differences. At both stages, the mid-height stem region was sampled to maintain a comparable positional context and to limit differences in tissue composition associated with the extensive xylem accumulation at the stem base (Figure 1B).

At 60 DAG, we found a substantial number of differentially expressed genes (DEGs) in both *ful2* and *stm3* (FDR < 0.05), with 641 DEGs shared between the two mutants (Figure 6A; Supplemental Data Set 1). This large overlap suggests that FUL2 and STM3 regulate a common set of genes, consistent with their reported ability to form a protein complex. The higher number of DEGs in the *stm3* stem may indicate a somewhat larger contribution of STM3 to transcriptional regulation, possibly because it can form homodimers/tetramers while FUL2 cannot (Jiang et al., 2022; Zahn et al., 2023). We next focused on the genes responding to the loss of both *FUL2* and *STM3* at 60 DAG. Notably, comparative analysis using PLAZA 5.1 assigned several of these shared genes to orthogroups containing Arabidopsis regulators of vascular development and secondary cell wall formation (Figure 6B) (Van Bel et al., 2022). *NAC14* (*Solyc06g073050*) and *NST1*-like (*Solyc03g080090*) belong to orthogroups containing NAC transcription factors associated with secondary growth, including NST1, a major regulator of secondary cell wall thickening in Arabidopsis (Mitsuda et al., 2005, 2007; Zhong et al., 2007). *Solyc10g055760* is a close homolog of NAC036, which has been associated with secondary cell wall formation and stem development (Kato et al., 2010; Porth et al., 2013).

**Figure 6.**
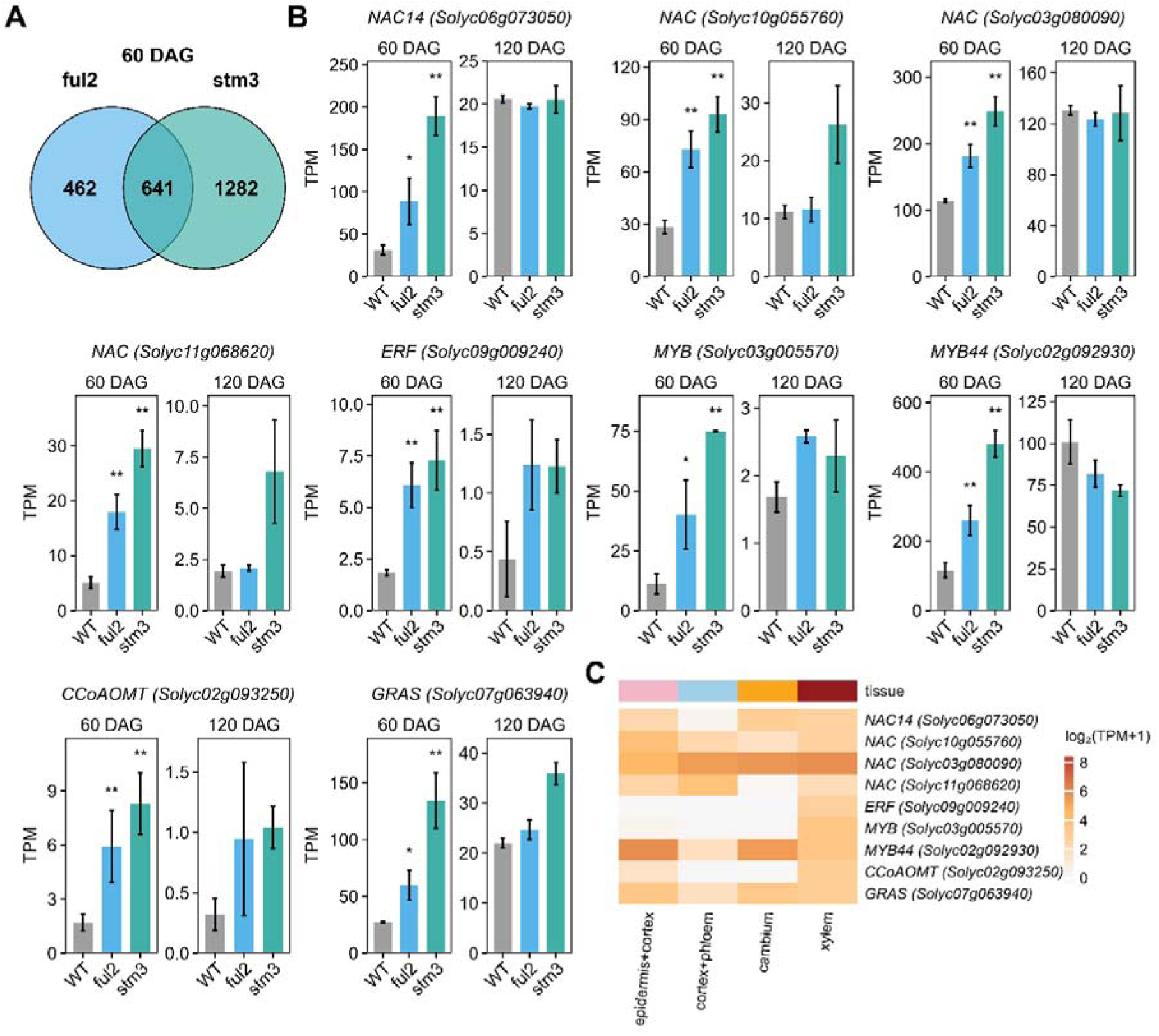
*FUL2* and *STM3* dependent gene expression changes at 60 DAG. **(A)** Venn diagram showing the overlap of differentially expressed genes (FDR < 0.05) in *ful2* and *stm3* compared with WT at 60 DAG. **(B)** Expression of genes selected from the intersection of DEGs shared between *ful2* and *stm3* at 60 DAG. Bars show mean TPM ± SE. **(C)** Spatial expression of the genes shown in (B) across the LCM defined stem tissues. The heatmap shows log_2_(TPM+1).

In addition to these NAC transcription factors, the DEGs include MYB-family transcription factors relevant to secondary growth. *Solyc03g005570* is orthologous to *MYB15*, which regulates lignin biosynthesis in Arabidopsis (Chezem et al., 2017), while *MYB44* (*Solyc02g092930*) groups with MYB factors including MYB117 and MYB3R1, associated with cell proliferation, meristem activity and developmental patterning (Gomez et al., 2011; Wang et al., 2018). Interesting are also the substantial upregulation of the ERF-family gene *Solyc09g009240* and GRAS family gene *Solyc07g063940,* of which close homologs are in Arabidopsis associated with increased stem biomass and tissue patterning, respectively (Karaba et al., 2007). In addition to these transcriptional regulators, the *CCoAOMT* gene, encoding caffeoyl-CoA 3-O-methyltransferase, an enzyme involved in monolignol biosynthesis, is also differentially expressed (Boerjan et al., 2003; Zhong et al., 2000).

We next examined the expression pattern of these genes in our LCM dataset and found that they are expressed in the vascular tissues, with in particular *CCoAOMT*, *MYB (Solyc03g005570)* and *ERF (Solyc09009240)* exhibiting xylem-specific expression (Figure 6C). To further examine their potential regulation by FUL2 and STM3, we searched the promoter regions of selected candidates for CArG-box motifs, which represent the binding sites of MADS-domain transcription factors. CArG-boxes were identified in the selected candidates (Supplemental Figure 8), suggesting that these genes may be directly regulated by FUL2-STM3. Thus, FUL2 and STM3 appear to repress genes related to vascular development and secondary wall formation at an early stage to regulate xylem differentiation, associated with the later secondary xylem mutant phenotype. Interestingly, the upregulation of these early targets in the mutants was much reduced, or absent, at 120 DAG.

### The FUL2 and STM3 dependent regulation differs at 120 DAG

To examine whether the transcriptional response to the loss of *FUL2* and *STM3* changes as stem development progresses, we analyzed the transcriptome of the mid-height stem region at 120 DAG. Interestingly, the candidates upregulated in the mutants at 60 DAG were no longer upregulated at 120 DAG, indicating a shift in the transcriptional response during stem development. At this stage, a much higher number of genes were commonly differentially expressed in *ful2* and *stm3* (Figure 7A; Supplemental Data Set 1). It is good to note that phenotypes at this stage are different between the mutants and the wild type, and that a substantial number of genes may be differentially regulated as a result of the shifts in tissue abundances.

**Figure 7.**
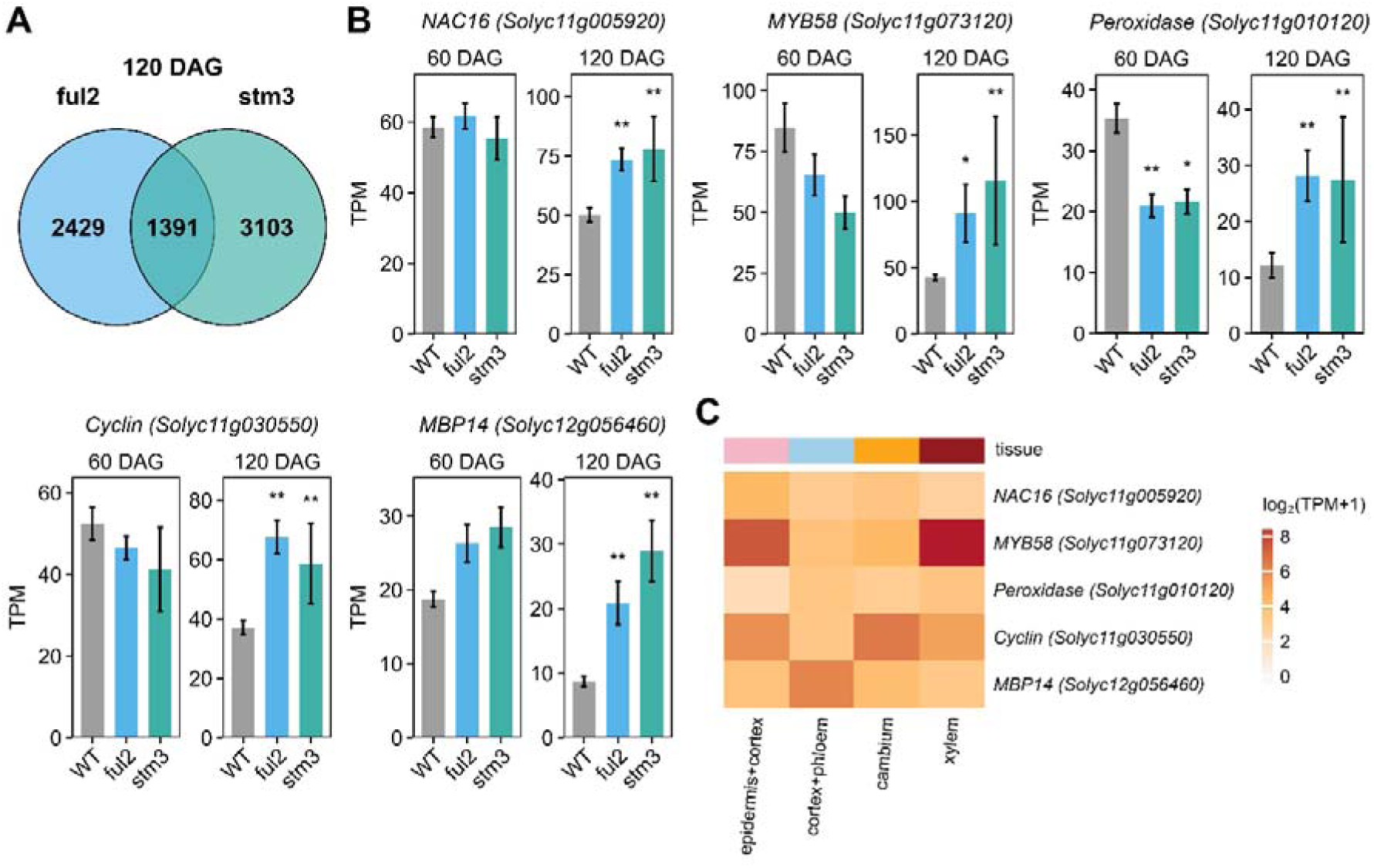
*FUL2* and *STM3* dependent regulation changes at 120 DAG. **(A)** Venn diagram showing the overlap of differentially expressed genes (FDR < 0.05) in *ful2* and *stm3* compared with WT at 120 DAG. **(B)** Expression of genes selected from the intersection of DEGs shared between *ful2* and *stm3* at 120 DAG. Bars show mean TPM ± SE. **(C)** Spatial expression of the genes shown in (B) across the LCM defined stem tissues. The heatmap shows log_2_(TPM+1).

The 120 DAG targets also include a *NAC* (*NAC16, Solyc11g005920)* and *MYB* TF genes (*MYB58, Solyc11g073120*) related to secondary cell wall formation, but these are different from those upregulated at 60 DAG (Figure 7B). *NAC16* (*Solyc11g005920*) is orthologous to Arabidopsis *NST* and *VND7,* which have established roles in secondary wall formation and vessel differentiation (Ailizati et al., 2021; Endo et al., 2015; Lehmann & Schneider, 2025; Yamaguchi et al., 2011). *MYB58* (*Solyc11g073120*) is orthologous to Arabidopsis *MYB58*, a key transcriptional activator of lignin biosynthesis and secondary cell wall formation (Zhou et al., 2009). The cyclin family gene *Solyc11g030550* was also upregulated, consistent with the established roles of cyclins in cell proliferation, cambial activity and secondary growth (Collins et al., 2015; de Oliveira et al., 2025; Shinozaki et al., 2018; Zheng et al., 2023). Interestingly, also *MBP14* (*Solyc12g056460*), a member of the broader tomato *SOC1*-like clade (Zahn et al., 2023), was upregulated in both mutants, suggesting a compensatory regulation within the *SOC1*-like family. In addition, *Solyc11g010120*, encoding a class III peroxidase probably participating in lignin polymerization and cell-wall modification (Fagerstedt et al., 2010; Marjamaa et al., 2009), was also upregulated. These genes were all expressed in the xylem fraction of the 60 DAG LCM data, although they did not show clear xylem-specific enrichment (Figure 7C). Their annotation and orthology to established regulators of xylem differentiation and secondary wall formation support a model in which FUL2 and STM3 regulate secondary growth mainly by repressing xylem differentiation and secondary cell wall formation.

### The FUL2 and STM3 dependent gene regulation depends on local secondary growth state

Next, we examined whether genes identified by RNA-seq at 60 and 120 DAG were associated with local secondary growth state. To this end, we checked their expression by qPCR at both mid-height and basal regions, including also the *quad ful soc1* mutant. We first analyzed 60 DAG candidates *NAC14*, *NAC (Solyc11g068620)* and *ERF (Solyc09g009240)*. Consistent with the RNA-seq, they were significantly upregulated in mid-height *ful2* and *stm3* stem segments. Surprisingly, the changes were weak or non-significant in *quad ful soc1* at the same stem position. Given that the extent of secondary growth highly depends on plant age, position in the stem, and possibly also the timing of the floral transition, we checked in different individuals of the same genotypes how stem phenotype varies around the mid height of the stem (Supplemental Figure 9A). Interestingly, we found that *quad ful soc1* stem diameter was only mildly enhanced compared to wild type at the 10^th^ internode, and narrower than *ful2* and *stm3* stems (Supplemental Figure 9B). We are not sure why *quad ful soc1* has a milder phenotype at mid-height, but it may be due to its much stronger delay in floral transition, whereas *ful2* and *stm3* show comparable flowering times (Supplemental Figure 9C). However, at the more developmentally advanced basal stem region, N*AC14, NAC* and *ERF* were also significantly upregulated in *quad ful soc1* (Figure 8A). This suggests that the weaker transcriptional response at mid-height may be associated with the local secondary growth state of the stem.

**Figure 8.**
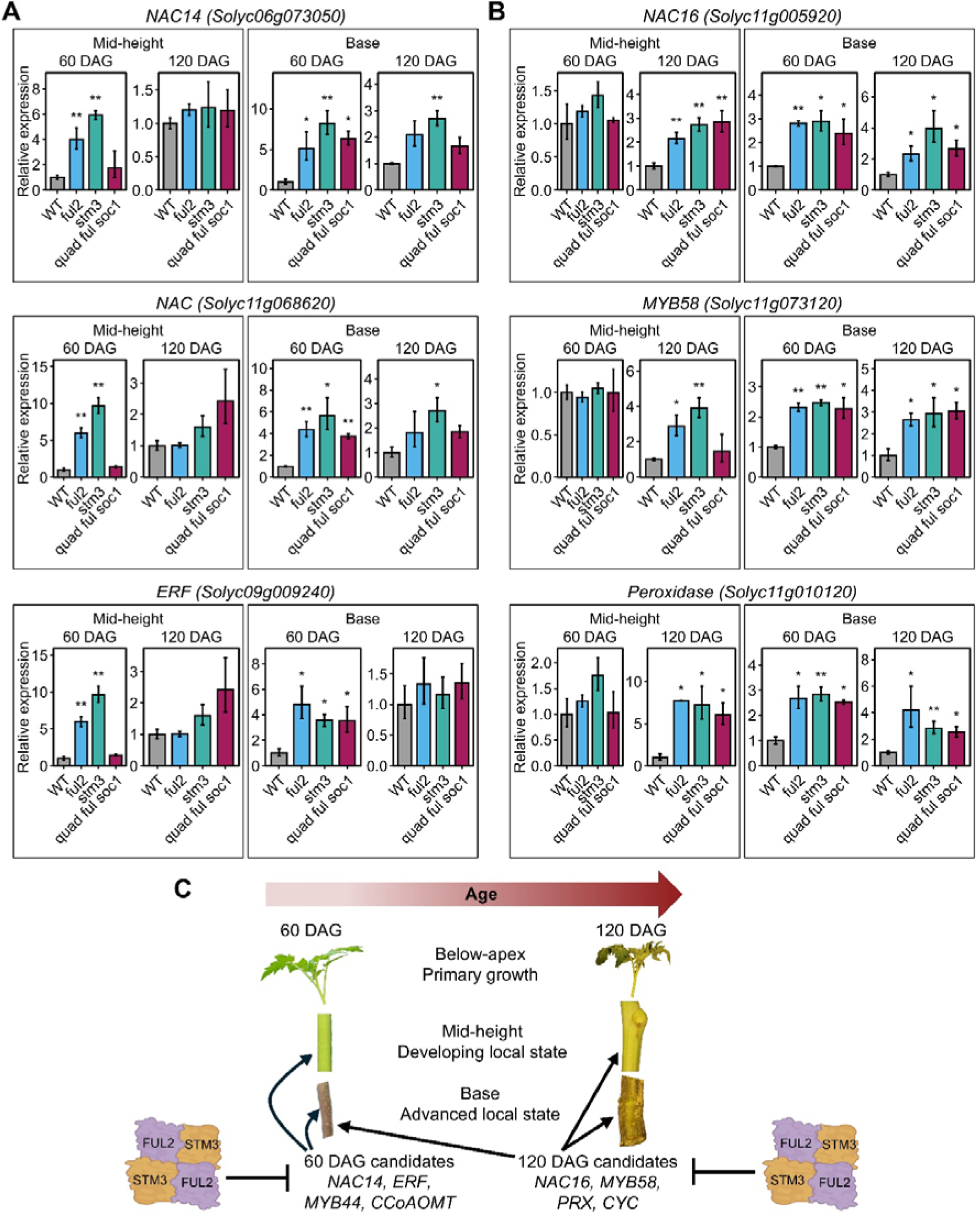
FUL2 and STM3 dependent regulation depends on local secondary growth state. **(A)** Relative transcript levels of the 60 DAG candidates in the mid-height and basal stem regions of 60 and 120 DAG WT, *ful2*, *stm3* and *quad ful soc1* plants. **(B)** Relative transcript levels of the 120 DAG candidates in the mid-height and basal stem regions of 60 and 120 DAG WT, *ful2*, *stm3* and *quad ful soc1* plants. Data are presented as mean ± SE. The asterisks indicate significant differences between each mutant and the corresponding WT at each age and stem position, determined using a two-tailed Student’s t-test at *p* < 0.05 (*) and *p* < 0.01 (**). **(C)** Schematic showing the relationship between local secondary growth state and the FUL2 and STM3 dependent transcriptional responses identified at 60 and 120 DAG. Genes identified as 60 DAG or 120 DAG candidates at the mid-height stem region can show altered expression at an earlier chronological age in the more developmentally advanced basal region.

We proceeded to examine *MYB58*, *NAC16* and the peroxidase *Solyc11g010120*, which were identified from the 120 DAG mid-height RNA-seq (Figure 8B). Interestingly, while we confirmed their differential expression at 120 DAG mid-height, significant upregulation was also already detected at the base of 60 DAG stems. Thus, genes identified at 120 DAG were already differentially expressed at 60 DAG in a stem region with a more advanced local secondary-growth state.

These results show that the transcriptional response identified by RNA-seq at 60 and 120 DAG mid-height samples does not reflect age alone. Instead, the effect of loss of FUL2 and STM3 on these genes is associated with the local secondary growth state of the stem, with responses detected earlier in the developmentally advanced basal region than at mid-height (Figure 8C).

## DISCUSSION

### FUL2 and STM3 have overlapping functions in suppressing secondary growth

Our results show that loss of either *FUL2* or *STM3* results in a pronounced increase in secondary xylem formation, identifying them as the major *FUL*- and *SOC1*-like genes suppressing secondary xylem development in tomato (Figure 4). The limited enhancement of the xylem phenotype in higher-order mutants indicates that the functions of *FUL2* and *STM3* are largely non-additive, and that they act together in the regulation of secondary growth. This relationship is particularly interesting in light of the molecular properties of *FUL2* and *STM3*. Both genes are also the key *FUL*- and *SOC1*-like regulators promoting the primary and sympodial transitions to flowering in tomato, where their encoded MADS-domain proteins interact in heterotetrameric complexes (Jiang et al., 2022, 2025b; Zahn et al., 2023). Unlike Arabidopsis FUL, which can form DNA-binding homodimers, tomato FUL2 does not homodimerize and therefore depends on interaction with other MADS-domain proteins for DNA binding (Jiang et al., 2022; Thoris et al., 2024). This difference in complex formation provides a possible explanation for the different genetic relationships observed between the two species. In Arabidopsis, FUL homodimerization could retain part of the regulatory activity in the absence of SOC1, consistent with the strong secondary growth phenotype observed in *ful soc1* double mutants only. In tomato, if STM3 is an important interaction partner of FUL2 in secondary vascular tissues, loss of either component could be sufficient to alter a shared regulatory activity, resulting in a similar phenotype of both *ful2* and *stm3*, and the limited further enhancement in higher-order mutants. Their overlapping expression and concordant transcriptional responses are consistent with this possibility.

Our finding should also be considered in relation to the previously reported effects of *SFT* and constitutive *FUL2* expression on secondary cell wall biogenesis in tomato (Shalit-Kaneh et al., 2019). These observations are not necessarily contradictory with our results because secondary xylem production and secondary wall biogenesis represent related but distinct developmental outputs. For a MADS-domain protein whose activity depends on interaction with other partners, elevated or ectopic expression could alter the composition or stoichiometry of transcriptional complexes and extend its activity beyond the spatial and developmental domains of the endogenous gene. The endogenous loss-of-function phenotypes we report here therefore provide information about FUL2 function in its normal developmental context that cannot be inferred directly from constitutive overexpression.

### FUL2 and STM3 dependent transcriptional responses depend on local secondary growth state

The enhanced secondary growth phenotype was evident at the later stage of stem development at 120 DAG. However, we observed the transcriptional response already at 60 DAG (Figure 6), before the anatomical phenotype was detected. At this earlier stage, *ful2* and *stm3* showed highly concordant responses involving NAC, MYB, ERF and GRAS transcription factors, together with genes associated with lignin and secondary cell wall metabolism. Many of these responses were not maintained at 120 DAG, when a largely distinct set of genes was affected, including regulators related to cell proliferation, vascular differentiation and secondary wall formation (Figure 7). The differences between the 60 and 120 DAG transcriptomes therefore indicate that loss of *FUL2* and *STM3* is associated with a changing transcriptional response as secondary growth progresses, rather than with a fixed set of direct and indirect downstream genes.

The temporal difference between these responses was further elucidated by examining the expression in the mutants at different positions along the stem (Figure 8). Genes identified from the 120 DAG mid-height transcriptome were already responsive at the more developmentally advanced base of 60 DAG plants, whereas the response of several 60 DAG genes in *quad ful soc1* was weak at mid-height but became significant in basal tissues. Thus, the transcriptional differences observed between 60 and 120 DAG cannot be explained by age alone. Instead, the effect of *FUL2* and *STM3* loss is associated with the local secondary growth state of the stem region.

Importantly, promoter reporter analyses further showed that the location of FUL2 and STM3 expression changes in the context of local secondary growth state. Both genes are active within secondary vascular tissues, with prominent activity in ray cells at later stages. Because rays form living radial connections across secondary vascular tissues and contribute to transport and communication between xylem and phloem (Pfautsch, Renard, et al., 2015; Spicer, 2014), this provides an interesting cellular context for their activity. Although the functional significance of ray expression remains unclear, our data suggest that the FUL2-STM3 complex may be required to repress secondary cell wall biogenesis and/or lignification to retain these living connections.

Altogether, we show that FUL2 and STM3 act within the secondary vascular tissues to suppress the progression of secondary xylem formation. Loss of either FUL2 or STM3 causes transcriptional responses involving NAC and MYB transcription factors, together with genes associated with secondary cell wall biogenesis. These responses change with the local secondary growth state, with different targets becoming apparent as secondary growth advances.

### The relationship between flowering time and secondary growth in tomato

Our tissue-resolved transcriptomic analysis showed conservation of regulatory programs controlling wood formation across dicot species, despite differences in growth habit and the extent and duration of secondary growth (Rodriguez et al., 2026). Interestingly, growth habit changes from herbaceous to (derived) woody are commonly accompanied by a delay in flowering time and increase in secondary growth, suggesting that these traits are coupled (Lens et al., 2012). Our LCM dataset provided an opportunity to examine if established regulators of the vegetative-to-reproductive transition are expressed within the stem tissues and may explain the link between both traits. The comparison revealed that, except for the xylem-specific expression of *FUL2*, there were no cambium- or xylem-associated flowering genes, which suggest that individual flowering regulators, rather than uniform activity of the flowering time regulatory network, can have additional functions during secondary growth. In addition to FUL2 and STM3, this may be the florigen SFT. Although our LCM data did not detect *SFT* transcripts in the stem, SFT protein may exert its effect on secondary growth as the mobile florigen, as indicated by the work of Shalit-Kaneh et al. (2019). In addition, other signals related to the floral transition, such as the gibberellin accumulation in Arabidopsis (Ragni et al., 2011; Sibout et al., 2008) may contribute to the connection between the timing of the floral transition and the extent of secondary growth. The fact that *FUL2* and *STM3* stem expression rises only after 30 DAG, when the floral transition has already occurred, could reflect broader age-dependent or developmental regulation and suggest that the upstream regulation is not completely linked to the floral transition. In Arabidopsis, the miR156–SPL age pathway can directly promote *FUL* expression, while SPL factors influence SOC1-dependent regulation (Hyun et al., 2016; Yamaguchi et al., 2009). Whether a related mechanism contributes to increase in *FUL2* or *STM3* expression in tomato stem remains unknown.

In conclusion, our results identify FUL2 and STM3 as negative regulators of secondary xylem formation in tomato. Their stem expression is not induced by the floral transition per se, but occurs somewhat later, and remains relatively stable at more advanced stages. At the same time, the downstream transcriptional response, which includes NAC and MYB TFs related to secondary xylem formation, varies with local secondary growth state.

## MATERIALS AND METHODS

### Plant materials and growing conditions

*Solanum lycopersicum* L. cv. Moneyberg was used in this study. Seeds were germinated on filter paper and transferred to rockwool blocks, where seedlings were grown for one week under natural light and supplied with 1 g/L Hyponex nutrient solution. Subsequently, plants were grown in a greenhouse under semi-controlled environmental conditions under a 16-h light/8-h photoperiod. Temperature was controlled using setpoints defined relative to sunrise and sunset, with values of 21°C and 16°C as maximum and minimum.

### Laser Capture Microdissection

Mid-height stem sections (1 cm) from sixty-day-old tomato plants were sampled and immediately fixed in Farmer’s fixative (75% ethanol, 25% acetic acid) ((Kerk et al., 2003). Samples were vacuum infiltrated twice for 15 minutes each and subsequently incubated at 4°C overnight. Following fixation, tissues were dehydrated through an ethanol series consisting of 75% and 96% ethanol (30 minutes each), followed by three changes of 100% ethanol (30 minutes each). Steedman’s wax was prepared by mixing polyethylene glycol 400 distearate and 1-hexadecanol in a 9:1 ratio, followed by melting and mixing at 65°C (Norenburg & Barrett, 1987; Steedman, 1957). Dehydrated samples in 100% ethanol were infiltrated in 1:1 mixture with Steedman’s wax at 37°C for 5 hours. This was followed by incubation in 100% Steedman’s wax at 37°C for 24 hours, during which the wax was refreshed three times. Samples were then transferred to a sterile petri dish and allowed to solidify. Embedded tissues were sectioned at a thickness of 12 µm using an automatic rotary microtome (Leica HistoCore AUTOCUT) and stretched on the nuclease-free water containing 40 µM DTT on 4 µm PEN-Membrane frame slides (Leica 11600289). Steedman’s wax was removed by washing the sections three times in 100% ethanol. Sections were air-dried and immediately subjected to laser microdissection (Leica LMD7000). Microdissected tissues were collected directly into extraction buffer from the Arcturus PicoPure RNA Isolation Kit (Thermo Fisher Scientific, Waltham, MA, USA). For each tissue type, eight sections were pooled per replicate. Total RNA was subsequently isolated using the same PicoPure kit according to the manufacturer’s instructions.

### RT-qPCR analysis

For RT-qPCR analysis, plants were grown as biological replicates. Stem samples were collected at the indicated DAG from defined positions below the apex, at mid-height, and at the stem base. Total RNA was extracted using the CTAB/LiCl method (Porebski et al., 1997), treated with TURBO DNase (Ambion, AM1907). cDNA was synthesized using SuperScript II Reverse Transcriptase (Invitrogen). RT-qPCR was performed using iQ SYBR Green Supermix (Bio-Rad) with a two-step amplification program of 40 cycles and an annealing temperature of 60°C. *ACTIN (ACT)* was used as the reference gene for normalization. Primer pairs previously described by Jiang et al. (2022) and Zahn et al. (2023) were used where applicable, and all primers are listed in Supplemental Data Set 1.

### GUS assay

The GFP-GUS reporter constructs were generated using the Golden Gate/MoClo cloning system (Weber et al., 2011). For the *STM3_pro_:GUS* the promoter of *STM3* was amplified from genomic DNA in five fragments. Fragment 4 was very repetitive and synthesized by GenScript. The amplified and synthesized fragments were cloned into the following level-0 GoldenGate vectors for modular assembly in a level-1 vector: pICH41276, pICH41308, pAGM1276, pAGM1263 and pICH41233. For the reporter construct, the level-1 vector was assembled in a level-2 vector together with a Golden Gate-compatible version of an eGFP-GUS module (Karimi et al., 2002), 35S::RFP and 35S::NPTII for selection of transgenic tissue. Stable transformation of tomato was performed using *Agrobacterium tumefaciens* strain C58C1 as described by van Roekel et al. (1993). Ploidy of regenerated plants was assessed by Iribov Analytical Services B.V. Diploid plants were used for reporter assay.

For histochemical GUS staining, stem samples were incubated in 50 mM sodium phosphate buffer (pH 7.0), 10 mM EDTA, 0.1% (v/v) Triton X-100, 2 mM potassium ferricyanide, 2 mM potassium ferrocyanide, and 2 mg mL⁻¹ 5-bromo-4-chloro-3-indolyl-β-D-glucuronic acid (X-Gluc). Samples were vacuum infiltrated twice for 5 min and incubated at 37°C until sufficient staining developed. Tissues were subsequently cleared in 70% ethanol to remove chlorophyll. Representative samples were sectioned and imaged.

### Sectioning, imaging and anatomical analysis

Freehand stem cross-sections were cut using a sharp razor blade. For paraffin embedding, approximately 1 cm stem segments were fixed in FAA and dehydrated through a graded ethanol series, followed by gradual replacement of ethanol with xylene. Samples were infiltrated with Paraplast Plus (Sigma Aldrich) at 60°C and embedded in fresh Paraplast Plus. Sections of 15 µm thickness were cut using a rotary microtome (Leica HistoCore AUTOCUT), floated on water, mounted on microscope slides, and air-dried. Sections were deparaffinized twice in 100% xylene for 10 min and rehydrated to 50% ethanol before staining.

Sections were stained with 1% Safranin O in 50% ethanol for 1 min, rinsed four times with MQ water, stained with 1% Alcian blue in 3% acetic acid (pH 2.5) for 3 min, and rinsed with MQ water. Sections were subsequently differentiated in 95% ethanol for 2 min and dehydrated in 100% ethanol and xylene. Stained sections were imaged using a Stemi 508 stereomicroscope equipped with an Axiocam 105 color camera (Zeiss). For confocal imaging of interfascicular xylem, stem sections were imaged using a Leica DM5500 confocal microscope (Leica Microsystems), with excitation at 488 nm and emission collected between 571 and 716 nm. Imaging settings were kept constant within each experiment. Anatomical measurements were performed using Fiji/ImageJ v1.52 (Schindelin et al., 2012).

### RNA-seq

RNA quality and concentration were assessed prior to library preparation using an Agilent 2100 Bioanalyzer. Low-input libraries for LCM RNA-seq were prepared by BGI Genomics, whereas standard RNA-seq libraries for bulk stem samples were prepared by Biomarker Technologies (BMK) GmbH. Both datasets were generated using 2 × 150-bp paired-end sequencing.

The raw reads were further processed to remove adapter sequences, low-quality reads, and contamination. Clean reads were then aligned to the tomato SL4.0 genome using HISAT2 (Kim et al., 2019) and counts were generated using featureCounts (Liao et al., 2014). Normalization and differential gene expression analyses were conducted using the DESeq2 package in R (Love et al., 2014). PCA was performed on the top 1,000 most variable genes using VST-transformed expression values generated by DESeq2.

### Identification of tissue specific, enriched and associated genes

Differential expression (DE) analysis was performed using the *DESeq2* package in R (Love et al., 2014), comparing each target tissue against all other tissues combined (Hou & Ji, 2024; Pullin & McCarthy, 2024). Log2 fold change values were shrinkage-adjusted to reduce noise in effect size estimates and improve the reliability of downstream analyses (Zhu et al., 2019). The enrichment ratio (ER) for each target tissue was calculated as the mean TPM in the that tissue divided by the maximum mean TPM among the remaining tissues, with a pseudocount of 0.1 added to the denominator. Genes were classified as tissue specific if they met the following criteria: FDR < 0.05, ER > 3, shrunken log2FC > 1, mean TPM in target tissue > 2, and detectable expression (raw counts ≥ 10) in at least two replicates of that tissue.

To define a broader set of tissue-enriched genes, less stringent filtering criteria were applied. Genes were classified as tissue-enriched if they satisfied the following thresholds: FDR < 0.05 and shrunken log2FC ≥ 1.5. In addition, tissue-associated genes were identified for a curated set of flowering time genes without applying a global statistical significance cutoff, and were instead defined based on an ER > 3. The epidermis+cortex depletion ratio (DR) was calculated as the maximum mean TPM across internal tissues (cambium, xylem, and phloem+cortex) divided by the mean TPM in epidermis+cortex, with a pseudocount of 0.1 added to the denominator. Expression heatmaps were generated using VST-transformed expression values, scaled per gene (row Z-score), and visualized using the *pheatmap* package in R.

### Enrichment analysis

Functional enrichment analysis was performed using gene set enrichment analysis (GSEA) (Subramanian et al., 2005). Genes were ranked based on the DESeq2 Wald statistic, and enrichment of MapMan functional categories was performed to obtain normalized enrichment scores (NES). Leading-edge genes were extracted and used for visualization. The hierarchical BINcodes of significantly enriched MapMan functional categories were manually curated, and category labels were simplified while preserving their functional interpretation. In addition, enrichment of selected gene sets (AspWood-derived gene sets) in the xylem tissue was evaluated using GSEA analysis. For GSEA, the same statistical approach as described above was applied. All genes included in the DE analysis were used as the background in this analysis.

### Orthology analysis

For cross-species comparative analysis, orthologs were identified using the PLAZA dicot workbench (Van Bel et al., 2022), requiring at least two evidence types: tree-based ortholog, orthologous gene family collinearity, or BHI family. Duplicate entries were removed to generate a set of tomato orthologs for downstream analyses.

### Lignin and structural carbohydrate quantification

The basal half of stems was dried at 48 °C and reduced to small particle sizes (between 1-2 mm) by means of a low-speed granulator. Extractives were then removed by sequential Soxhlet extraction with toluene, ethanol, and water, following the procedure described by Appiah et al. (2026) with modifications based on NREL/TP-510-42619 (Sluiter et al., 2008). Extractive-free material was further milled to a 0.2–0.8 mm particle size range and homogenized by compositing samples from the top, middle and bottom layers of the bulk extractive-free material. The final composite was subjected to two-step acid hydrolysis according to NREL/TP-510-42618, as described by Appiah et al. (2026), with additional steps for protein content correction. In details, samples were hydrolyzed in 72% (w/w) H₂SO₄ at 30 °C for 1 h, followed by dilution to 4% H₂SO₄ and hydrolysis at 121°C for 1 h. Acid-insoluble lignin was recovered by vacuum filtration and determined gravimetrically after correction for ash and protein, whereas acid-soluble lignin in the filtrate was quantified by UV–Vis spectrophotometry. Total lignin content was calculated as the sum of acid-insoluble and acid-soluble lignin. Monomeric sugars released during hydrolysis were quantified by HPLC using an Aminex HPX-87H column and converted to structural carbohydrate content using the appropriate anhydro correction factors. Total structural carbohydrate content was calculated as the sum of the quantified structural carbohydrates. Lignin and carbohydrate contents were reported on extractives-free dry-weight basis. An overview of the complete composition including extractives is provided in Supplementary Figures 3C-G.

### Accession Numbers

*Solanum lycopersicum* gene identifiers (Solyc IDs) for the genes referenced in this article can be found in Supplemental Data Set 1. The raw RNA-seq data generated in this study are available at the European Nucleotide Archive (http://www.ebi.ac.uk/ena/) under study accessions PRJEB129214 (bulk RNA-seq of f*ul2*, *stm3* and WT) and PRJEB129245 (LCM RNA-seq).

## Supporting information

Supplemental Data Set

## ACKNOWLEDGMENTS

We thank Remko Offringa for reading the manuscript and providing valuable feedback. We thank Indira Sanchez Bretones and Chris Roelofsen for their help in generating the STM3 reporter line. We thank Erik Limpens for demonstrating the operation of laser capture microdissection system, and the Unifarm staff for their help in caring for tomato plants and collecting seeds in the greenhouse. We also thank all members of the WOODLIG consortium for the constructive discussions. This work was supported by the Dutch Research Council (NWO; grant OCENW.XL21.XL21.059).

## AUTHOR CONTRIBUTIONS

M.B. conceived the project and designed the experiments; K.P. performed phenotyping, LCM, RNA-seq experiments, and qPCR analyses; A.P.A. performed biochemical analyses of lignin and carbohydrates; I.E.Z. generated the STM3 reporter line; E.B. sectioned and imaged mid-height stem regions from 120 DAG samples; R.v.V. performed the phylogenetic analysis; P.J.D., G.C.A. and M.B. supervised the project; M.B. and K.P. analyzed the data, prepared the figures, and wrote the manuscript. All authors read and approved the final version of the manuscript.

