## Supplemental information for "Secondary growth in the tomato stem is suppressed by the flowering regulators FUL2 and STM3"

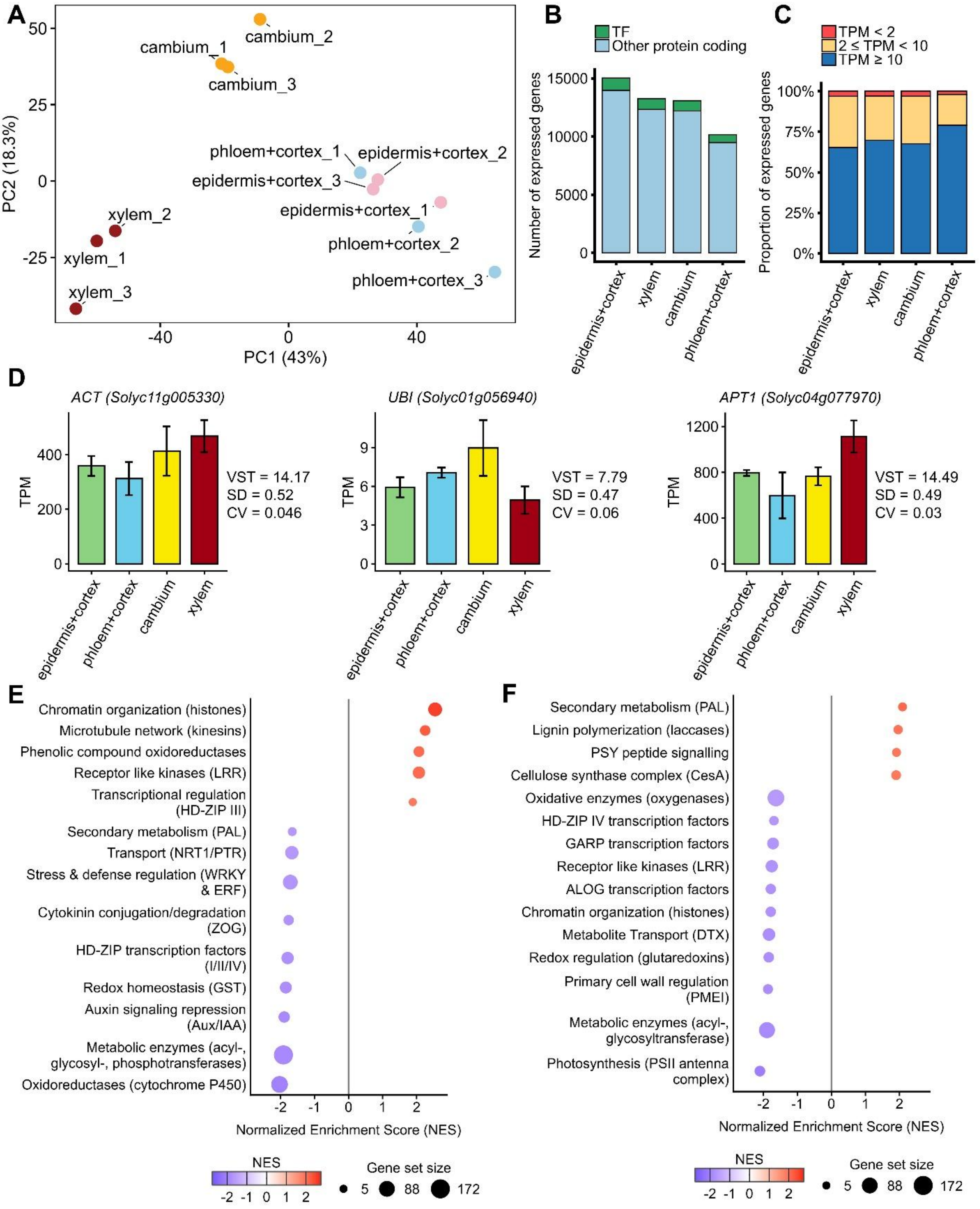


**Supplemental Figure 1. Expression profiles and quality assessment of samples analyzed by LCM RNA-seq. (A)** PCA of VST normalized read counts. **(B)** Number of expressed TFs and non-TF genes detected in each tissue type. **(C)** Proportion of genes within different expression-level categories based on TPM. **(D)** Expression of housekeeping genes *ACT*, *UBI*, and *APT1*. Bars represent mean TPM values ± SEM. SD, standard deviation; CV, coefficient of variation. **(E)** MapMan-based GSEA of cambium DEGs displaying normalized enrichment scores (NES) for significantly enriched and depleted functional categories. **(F)** MapMan-based GSEA of xylem DEGs showing NES for significantly enriched and depleted functional categories.

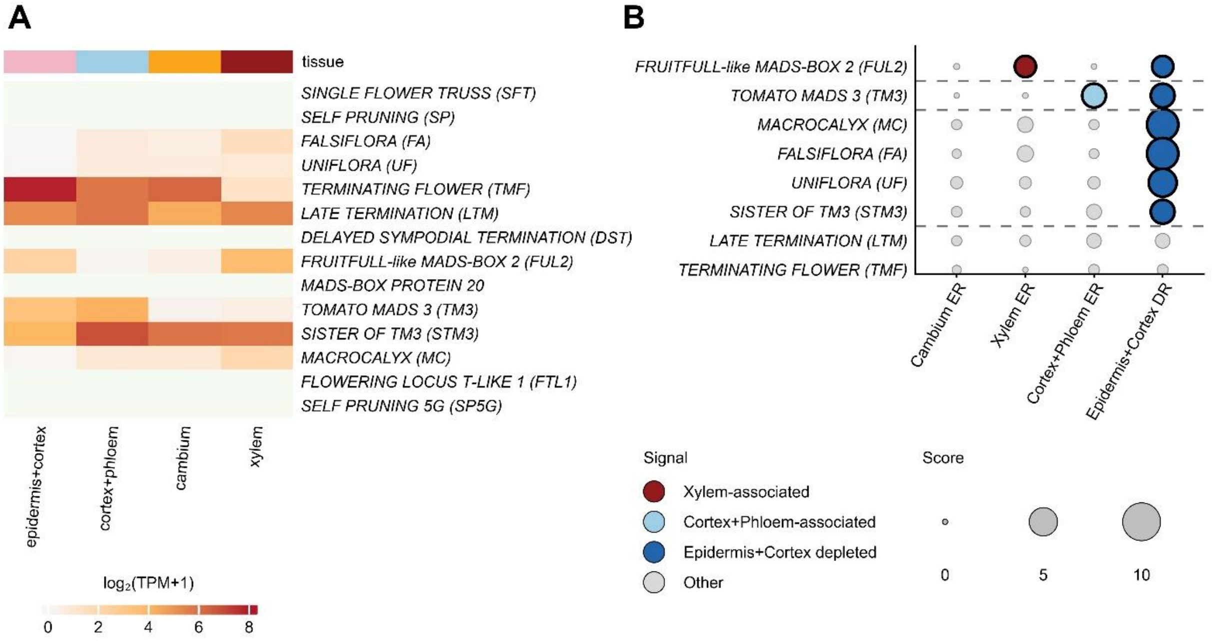


**Supplemental Figure 2. Expression profiles of genetically established regulators of vegetative to reproductive transition across tomato stem tissues.** **(A)** Heatmap showing expression of genetically established key regulators of primary and/or sympodial vegetative to reproductive transition across LCM samples. **(B)** Tissue association of genetically established key regulators of vegetative to reproductive transition in tomato detected in LCM samples (TPM ≥ 1). Gene with ER>3 were classified as preferentially associated with that tissue, whereas genes with depletion ratio (DR) > 3 in epidermis+cortex sample were classified as preferentially associated with inner vascular tissues.

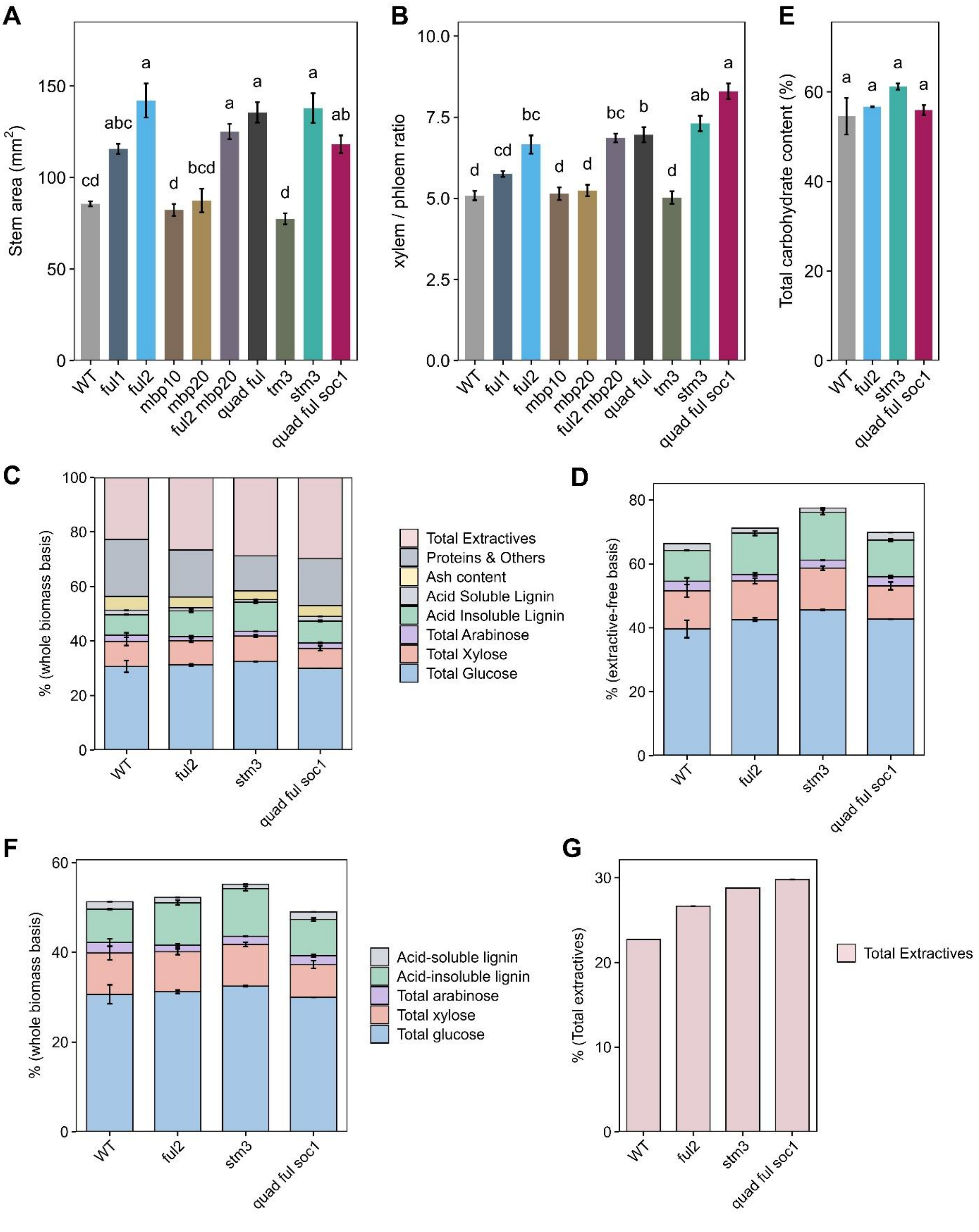


**Supplemental Figure 3. Quantification of stem area, xylem-to-phloem ratio, and biomass composition in 120 DAG stems.** **(A)** Stem area. **(B)** Xylem-to-phloem ratio. **(C)** Biomass composition of the bottom half of the stem on a whole-biomass basis. **(D)** Lignin and carbohydrate composition on an extractive-free biomass basis. **(E)** Total carbohydrate content on an extractive-free biomass basis. **(F)** Lignin and carbohydrate composition on a whole-biomass basis. **(G)** Total extractives content on a whole-biomass basis. Statistical differences were determined by one-way ANOVA followed by Tukey's HSD test ($p < 0.01$). Different letters indicate statistically significant differences. Values are shown as mean ± SE.

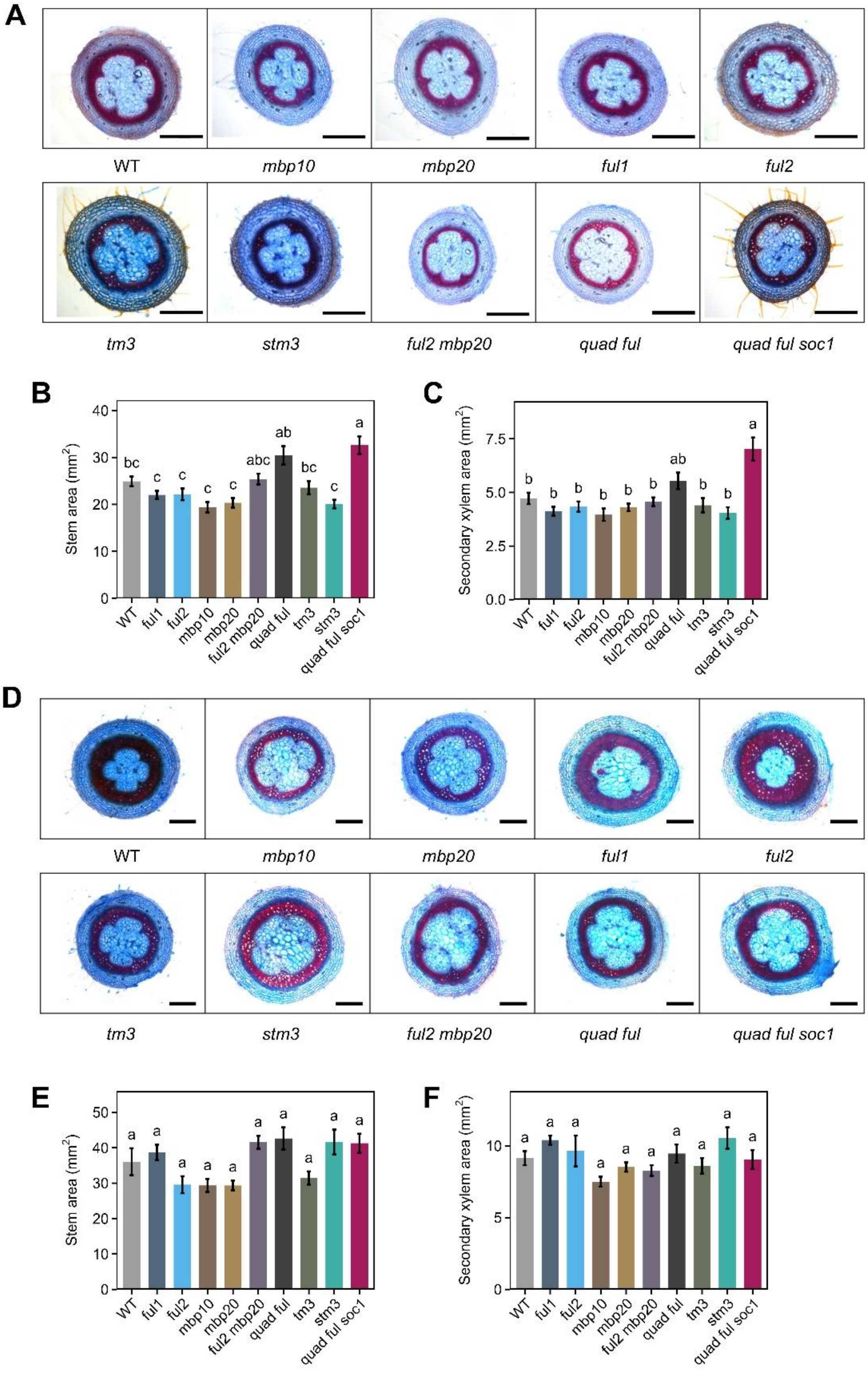


**Supplemental Figure 4. Secondary growth phenotypes of mutants of *FUL*-like and *SOC1*-like genes in the basal stem at 30 and 60 DAG. (A-C)** Secondary growth phenotypes at 30 DAG in the basal internode. Among the tested mutants, only *quad ful soc1* showed a mild increase in stem area and secondary xylem area compared with WT. **(D-F)** Secondary growth phenotypes at 60 DAG in the basal internode. At this stage, all mutants showed stem and secondary xylem areas comparable to WT. Statistical differences were determined by one-way ANOVA followed by Tukey's HSD test ($p$ < 0.01). Values are shown as mean ± SE (n = 5). Different letters indicate statistically significant differences.

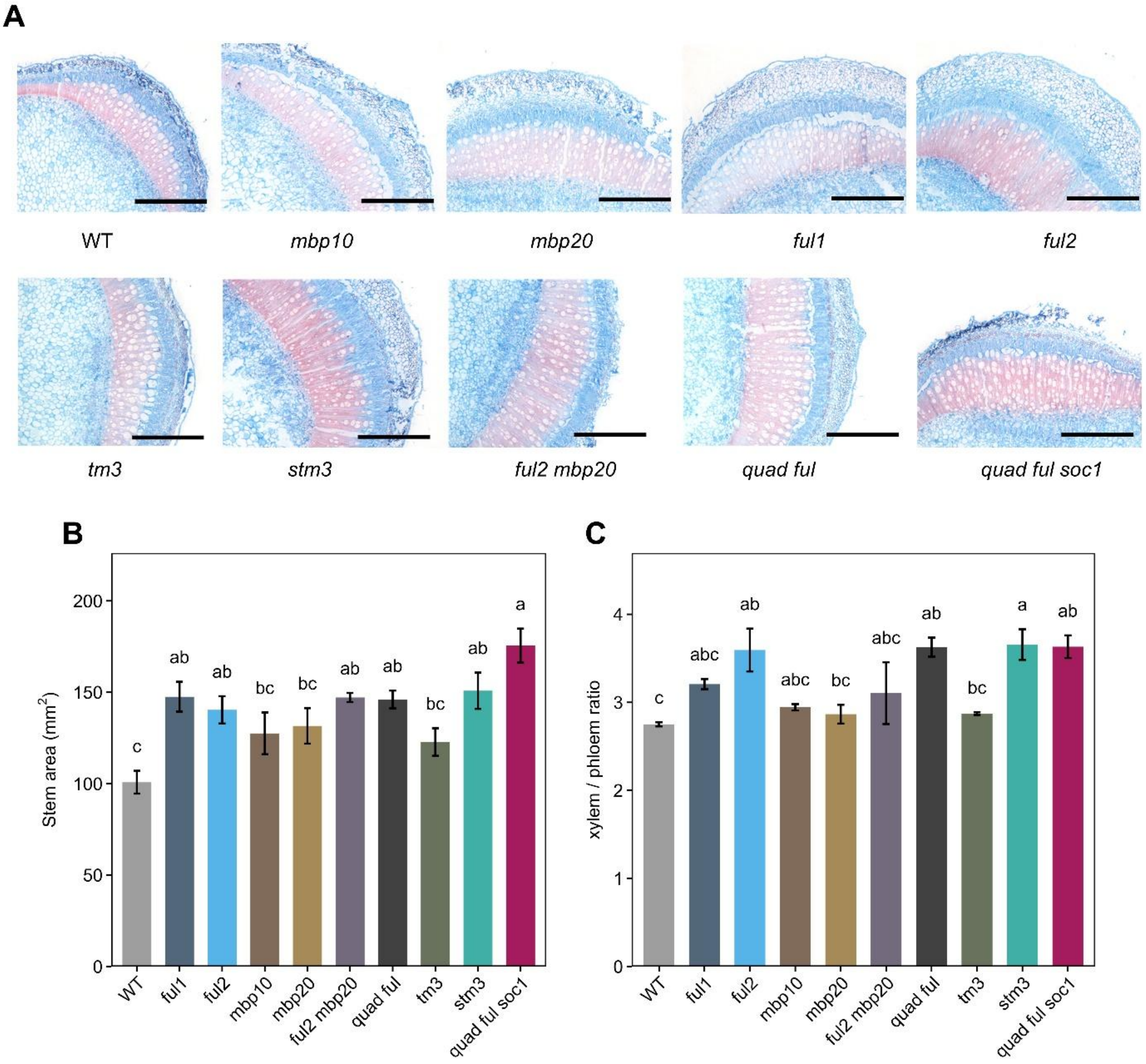


**Supplemental Figure 5. Secondary growth phenotypes of *FUL*-like and *SOC1*-like mutants in the mid-height stem region at 120 DAG. (A)** Representative cross-sections from the mid-height stem region of the indicated mutants. Scale bars, 2 mm. **(B and C)** Stem area (B) and xylem-to-phloem ratio (C) at 120 DAG. *ful2*, *stm3*, and *quad ful soc1* showed significant increases in both stem area and xylem-to-phloem ratio compared with WT. Statistical significance was assessed by one-way ANOVA followed by Tukey's HSD test ($p < 0.01$). Different letters indicate statistically significant differences. Values are shown as mean ± SE (n = 4).

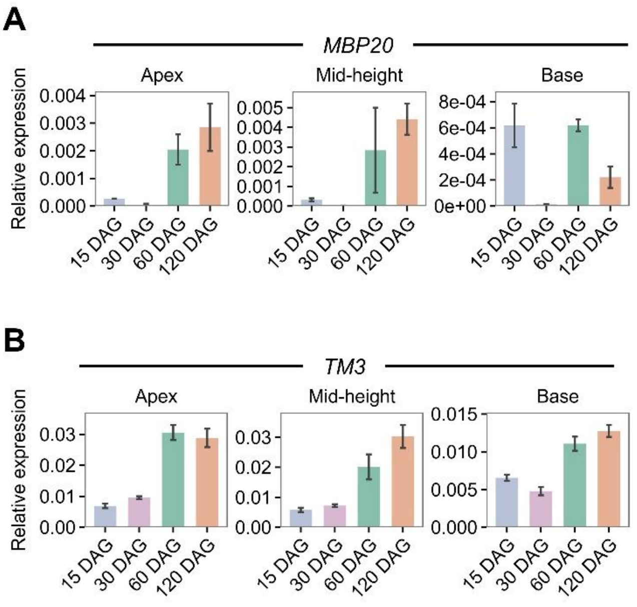


**Supplemental Figure 6. Expression patterns of *MBP20* and *TM3* across the tomato stem.** *TM3* showed a expression pattern similar to that of *FUL2* and *STM3*, and *MBP20* was expressed at comparatively low levels.

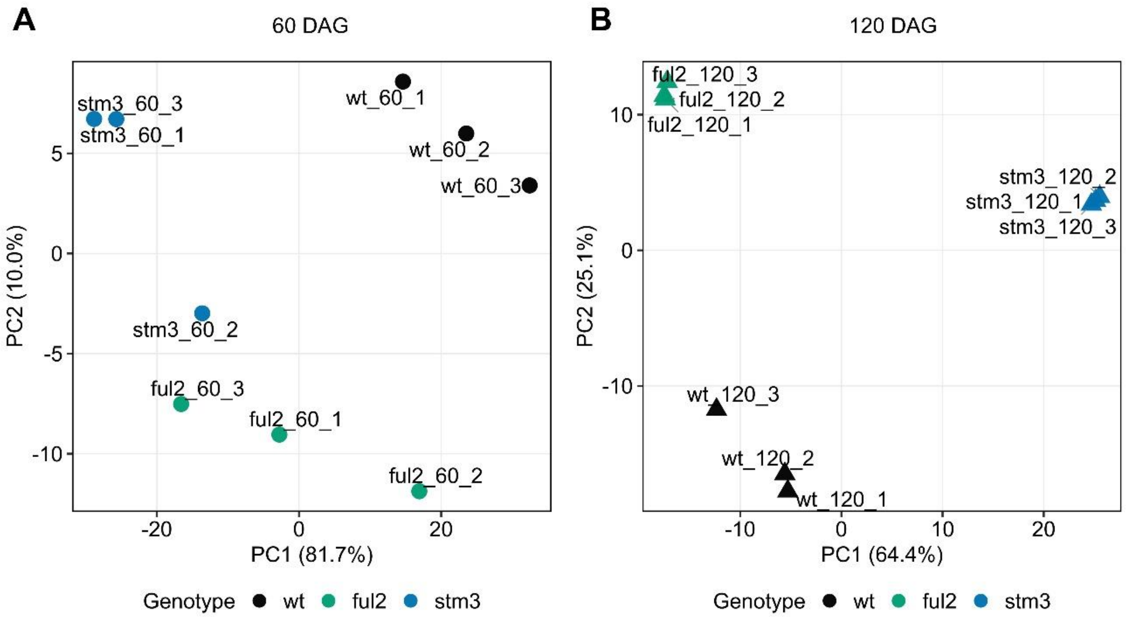


**Supplemental Figure 7. PCA of *ful2* and *stm3* at 60 DAG (A) and 120 DAG (B)**

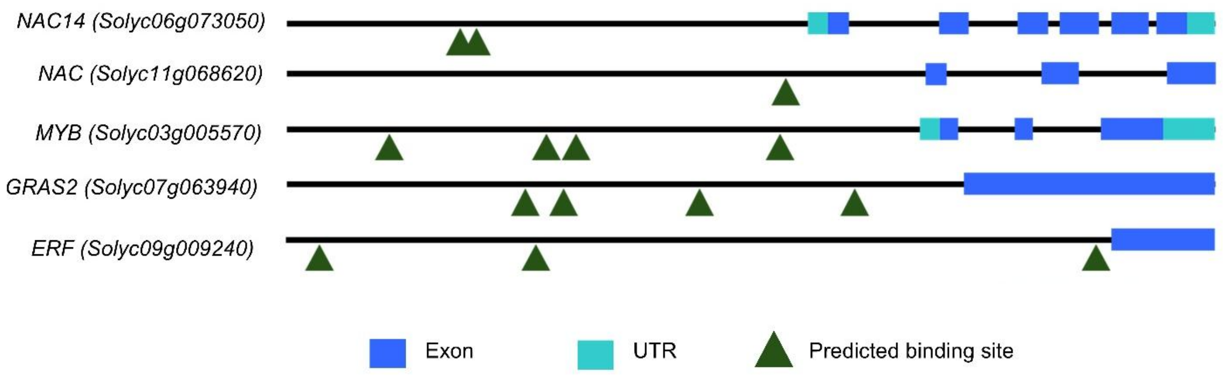

**Supplemental Figure 8. Predicted CArG-box motifs in downstream candidates of FUL2 and STM3.** Schematic representation of the 5-kb upstream regions and gene structures of selected downstream candidates. Green triangles indicate predicted CArG-box motifs. Blue boxes indicate exons and cyan boxes indicate UTRs.

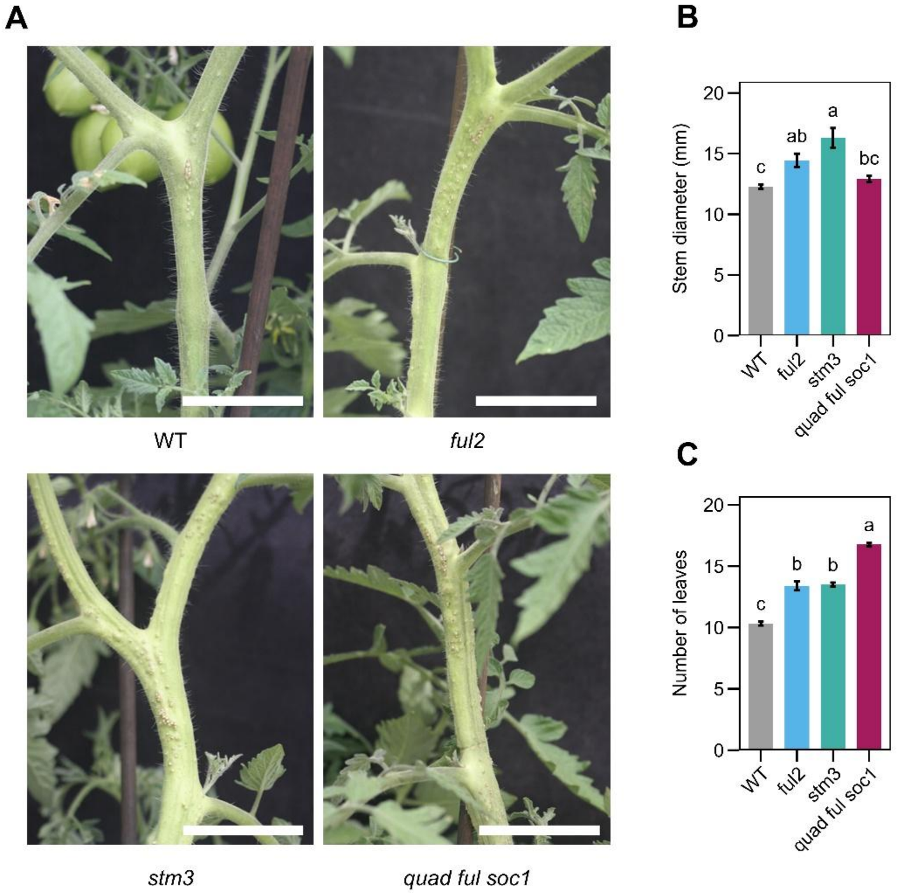


**Supplemental Figure 9. Flowering time and radial stem growth in *ful2, stm3* and *quad ful soc1* at 75 DAG. (A)** Representative stems of 75 DAG WT, *ful2*, *stm3* and *quad ful soc1* plants showing the tenth internode used for stem diameter measurements. This position corresponds to the internode immediately below the first inflorescence in WT. Scale bars, 5 cm. **(B)** Stem diameter at the tenth internode in WT and mutant plants at 75 DAG. **(C)** Number of leaves produced before the first inflorescence in WT, *ful2*, *stm3* and *quad ful soc1* plants. Statistical differences were determined by one-way ANOVA followed by Tukey's HSD test ($p$ < 0.01). Different letters indicate statistically significant differences. Values are shown as mean ± SE.

Supplemental Table 1. Curated tomato regulators of the vegetative to reproductive transition with genetic evidence for function.

| Tomato gene | Gene ID | Arabidopsis homolog / related gene family | Transition context | Role in reproductive transition | Genetic evidence | References |
|---|---|---|---|---|---|---|
| *SINGLE FLOWER TRUSS (SFT)* | *Solyc03g063100* | *FLOWERING LOCUS T (FT)* | Primary + sympodial | Major florigen that promotes reproductive transition and flowering | *sft* mutants flower late | Lifschitz et al., 2006; Molinero-Rosales et al., 2004 |
| *SELF PRUNING (SP)* | *Solyc06g074350* | *TERMINAL FLOWER1 (TFL1)*-like PEBP | Mainly Sympodial | Maintains the vegetative phase of sympodial meristems | *sp* mutants show progressive decrease in leaf number per sympodial unit, resulting in determinate shoot | Pnueli et al., 1998 |
| *FALSIFLORA (FA)* | *Solyc03g118160* | *LEAFY (LFY)* | Primary + sympodial | Promotes flowering and floral meristem identity | *fa mutants* show delayed flowering in both the primary shoot and early sympodial shoots | Molinero-Rosales et al., 1999, 2004 |
| *UNIFLORA (UF)* | *Solyc09g005070* | bHLH family | Primary + sympodial | Regulates meristem reprogramming during floral transition and inflorescence meristem initiation | *uf* mutants are late flowering, produce solitary flowers, and flowering delay is season dependent | Dielen et al., 2004; Meir et al., 2021 |
| *TERMINATING FLOWER (TMF)* | *Solyc09g090180* | ALOG family | Mainly Primary | Represses meristem maturation and floral specification during primary transition | *tmf* mutants show early flowering and the primary inflorescence converts into a solitary flower | MacAlister et al., 2012 |
| *LATE TERMINATION (LTM)* | *Solyc01g100600* | Kelch-repeat protein | Primary | Coordinates SAM maturation with the timing of reproductive transition | *ltm* mutants are severely late flowering despite precocious doming of the vegetative SAM | Tal et al., 2017 |
| *DELAYED SYMPODIAL TERMINATION (DST)* | *Solyc03g006900* | MYB-SANT-like DNA-binding protein with a kinase domain | Primary + sympodial | Promotes floral transition through a florigen independent pathway acting upstream of *FA* | Loss of *DST* delays flowering, and combined loss of *DST* and *SFT* produces an essentially ever-vegetative phenotype | Meir et al., 2021 |

| | | | | | | |
|---|---|---|---|---|---|---|
| *FRUITFULL-like MADS-BOX 2 (FUL2)* | *Solyc03g114830* | *FUL*-like | Primary + sympodial | Promotes vegetative to reproductive transition | Loss of *FUL2* delays primary and sympodial flowering | Jiang et al., 2022 |
| *MADS-BOX PROTEIN 20* | *Solyc02g089210* | *FUL*-like | Primary + sympodial | Promotes vegetative to reproductive transition | Loss of *MBP20* delays primary and sympodial flowering | Jiang et al., 2022 |
| *TOMATO MADS 3 (TM3)* | *Solyc01g093965* | *SOC1*-like | Primary + sympodial | Promotes vegetative to reproductive transition | Loss of TM3 causes delayed flowering in primary and sympodial shoots | Zahn et al., 2023 |
| *SISTER OF TM3 (STM3)* | *Solyc01g092950* | *SOC1*-like | Primary + sympodial | Promotes vegetative to reproductive transition | Loss of STM3 causes delayed flowering in primary and sympodial shoots | Zahn et al., 2023 |
| *MACROCALYX (MC)* | *Solyc05g056620* | *APETALA1 (AP1)* | Primary + sympodial | Promotes flowering redundantly with *FUL2* and *MBP20* | *mc* has a modest primary flowering effect but strongly enhances *ful2 mbp20*; loss of all three causes extremely delayed sympodial flowering | Jiang et al., 2025 |
| *FLOWERING LOCUS T-LIKE 1 (FTL1)* | *Solyc11g008650/ Solyc11g008660* | *FT-like* | Primary | Promotes reproductive transition under short-day conditions | Loss of *FTL1* delays flowering under short days; natural variation at *FTL1*, together with *SP5G*, contributed to the evolution of day-neutral flowering in cultivated tomato | Song et al., 2020 |
| *SELF PRUNING 5G (SP5G)* | *Solyc05g053850* | *FT*-like | Primary | Represses floral transition under long day conditions | *sp5g* mutants are early flowering under long days | Soyk et al., 2017 |
